# OTTER: Orchestrated Transcriptomic, Tumor-xenograft (PDX/CDX), and Epigenomic Reporting Workflow

**DOI:** 10.64898/2026.10.24.761506

**Authors:** Yanhua Zheng, Xin Liu, Yingyu Yi, Xiaojun Sun, Fang Liu, Xiaojing Yan, Cheuk-Him Man, Hongcang Gu, Xiaoxue Wang, Fan Zhang

## Abstract

**Summary:** Tumor xenograft sequencing libraries inevitably contain infiltrating murine host reads, but conventional deconvolution tools fail on bisulfite sequencing (BS-seq) because chemical C-to-T deamination collapses sequence entropy and destroys graft recall (*<* 0.015%). We present OTTER, an automated, compiled multi-omics workflow framework that decouples execution into static domain operators, an immutable reference registry, and a native strand-aware edit-distance engine, restoring bisulfite deconvolution (52%–54% recall across a 0%–99% compositional gradient) while preserving complete decision parity on transcriptomics.

**Availability and implementation:** OTTER is implemented in Go and Rust, runs on x86 64 Linux (including CentOS 7 with glibc 2.17), and is freely available open-source under the MIT License at https://github.com/otterlab-bio/otter; test and validation data are provided at https://huggingface.co/datasets/fallingstar10/otter-data.

**Supplementary information:** Supplementary data are available with this preprint.

## Introduction

Tumor xenografts, primarily patient-derived xenograft (PDX) and cell line-derived xenograft (CDX) models (collectively designated XDX models), represent a cornerstone of translational cancer biology, functional genomics, and preclinical drug evaluation (Woo et al., 2019). By engrafting human tumor biopsies or characterized cancer cell lines into immunodeficient mice, XDX models preserve cellular heterogeneity, histological architecture, and therapeutic responsiveness far more effectively than traditional in vitro two-dimensional cultures.

A fundamental computational challenge in xenograft profiling is the infiltration of murine host stroma (comprising fibroblasts, immune cells, endothelial cells, and extracellular matrix) into the human graft: xenograft sequencing libraries are invariably chimeric mixtures of human graft and mouse host reads, with murine content commonly ranging from 10% to over 70% depending on tumor type, passage number, and vascularization (Jo et al., 2019; Woo et al., 2019). Unless these host reads are removed computationally, they cause severe downstream artifacts, including false somatic variant calls, distorted gene-expression counts, and skewed DNA methylation profiles.

Several tools have been developed to separate host from graft reads. Early strategies such as Xenome (Conway et al., 2012) use *k*-mer-based taxonomic classification, trading base-level precision for speed; alignment-based utilities such as Disambiguate (Ahdesmäki et al., 2016) and XenofilteR (Kluin et al., 2018) compare alignment scores and edit distances across dual-species alignments; and pseudo-alignment tools such as Xengsort (Zentgraf and Rahmann, 2021) provide fast, memory-efficient transcriptomic read sorting.

These methods, however, share two critical limitations. First, they are fundamentally inapplicable to bisulfite sequencing (BS-seq, including RRBS and WGBS): sodium bisulfite deaminates unmethylated cytosines to uracils (amplified as thymines), contracting the four-letter genomic alphabet to an asymmetric three-letter code, so standard aligners and edit-distance discriminators misinterpret genuine conversions as mismatches, causing cross-species mapping failure and spurious methylation profiles (Krueger and Andrews, 2011). Second, existing tools are fragmented point solutions without multi-omics integration (Bhandari et al., 2025), executed as fragile scripts lacking automated quality control, cluster scheduling, and checkpointed recovery.

To resolve these challenges, we present **OTTER** (https://github.com/otterlab-bio/otter), an automated, production-scale multi-omics workflow framework that executes end-to-end preclinical profiling across RRBS, WGBS, RNA-seq, and matched PDX/CDX models, spanning from raw FastQ quality control and dual-species alignment through bisulfite-aware host-read deconvolution, mate-pair resynchronization, expression quantification, alternative splicing, and genome-wide CpG methylation-matrix construction. We validate OTTER on two production supercomputing clusters and authentic public SRA cohorts (a 32.1M-fragment RNA-PDX and a 45.2M-fragment BS-PDX accession), on dual 0%–99% compositional gradients, and through factorial parameter audits.

## System and methods

### The xdxtools prototype

Our initial prototype, xdxtools (https://github.com/Genomiclab/xdxtools), was an R package (R *≥* 3.5) built around Snakemake (Mölder et al., 2025) workflows that invoke FastQC, Trim Galore, Samtools, Bismark (Krueger and Andrews, 2011), STAR (Dobin et al., 2013), Picard, HTSeq (Anders et al., 2015), XenofilteR (Kluin et al., 2018), and rMATS (Shen et al., 2014) for RRBS, WGBS, and RNA-seq XDX scenarios, coordinated by an R6 controller (BeaverGandalf; Supplementary Note 1). Its foundational innovation was extending XenofilteR deconvolution to bisulfite-aware alignments via Picard tag recalibration (`IS_BISULFITE_SEQUENCE=true`), providing end-to-end processing from raw FastQ files to analysis-ready Bioconductor objects (`methrix` (Mayakonda et al., 2020) and bsseq (Hansen et al., 2012)) with fault-tolerant execution across diverse computing infrastructures. However, enterprise cluster deployments unmasked three systemic systems-level bottlenecks: (1) multi-omics Conda dependency graphs frequently triggered SAT-solver deadlocks and multi-gigabyte storage bloat; (2) institutional clusters anchored to legacy Linux distributions (CentOS 7, glibc *≤* 2.17) failed with dynamic-linking symbol errors (`GLIBC_2.XX not found`), presenting an insurmountable barrier for non-root researchers; and (3) Snakemake’s shared-filesystem lock protocol suffered severe lock contention and false-positive aborts on parallel Lustre/NFS storage under massive task parallelism. These physical constraints fundamentally motivated a ground-up architectural overhaul into OTTER.

### The OTTER architecture

To eliminate these bottlenecks, OTTER decouples workflow execution into explicit compiled layers (Fig. 1): (1) the core toolchain ships as statically compiled Go and Rust binaries, eliminating runtime R dependencies and glibc symbol drift; (2) a lightweight Rust environment manager (`enva`) manages minimal Conda-compatible environments so that only upstream heavy aligners (Bismark, Bowtie2, STAR) require resolved scientific environments; and (3) an embedded Go orchestrator (`craftmake`) coordinates asynchronous task execution via SQLite state persistence, completely eliminating network-filesystem lock contention (Supplementary Note 2, Sections S2.1–S2.4; Supplementary Note 4, Section S4.2). Reference genomic assets are cryptographically sealed in an immutable Reference Genome Registry (`$OTTER_REFERENCE_ROOT`), while canonical execution snapshots bind inputs, configurations, and toolchain manifests into versioned contracts (otter.run/v1; Supplementary Note 2, Section S2.5).

**Figure 1.**
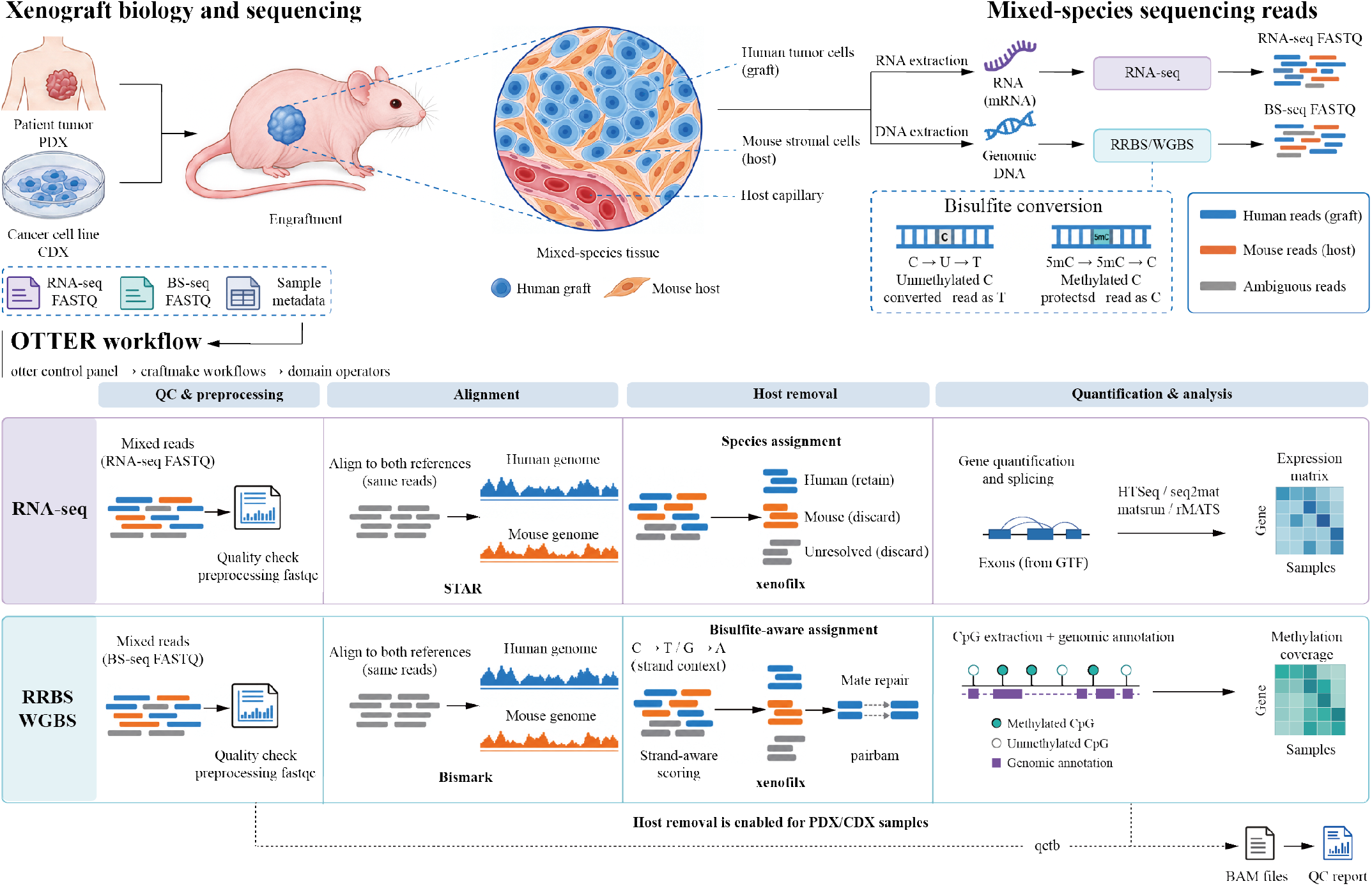
The decoupled OTTER system architecture. The otter Go control plane and immutable Reference Registry feed the `craftmake` orchestrator, which dispatches statically compiled domain operators (`fastqcx, xenofilx, methx, pairbam, seq2mat, matsrun, qctb`) and the `bamdriver` primitive library, `enva` isolates only the heavyweight aligners, and outputs are published under versioned contracts.

### Compiled domain operators

Seven static domain operators replace the prototype’s legacy R scripts across each computational bottleneck:

- `fastqcx` merges the core quality-control metrics of FastQC and seqkit into a single pass, producing output numerically identical to the combined FastQC + SeqKit oracle on identical inputs (Supplementary Note 4, Section S4.3);
- `xenofilx` reimplements the Picard + XenofilteR flow on our Go `bamdriver` library with native strand-aware NM recalculation, eliminating Java and `samtools` system dependencies while achieving 100% decision parity on RNA-PDX and, on bisulfite data, +0.63%–0.70% higher graft recall than the prototype across a 0%–99% BS-seq gradient (rescuing *≈* 4,000 additional authentic human fragments per million reads at 60% human content; Supplementary Note 4, Section S4.5);
- `methx` replaces the `methrix` R package (Mayakonda et al., 2020) in Bismark downstream processing, extracting CpG loci directly from FASTA and optionally annotating them against GTF; evaluated against Methrix’s canonical BSgenome workflow on full-genome eRRBS data, `methx` achieves a 2.41-fold speedup (1 m 32 s vs 3 m 41 s) and slashes peak RSS from 18.62 GiB to 6.42 GiB (a 65.5% memory reduction), emitting versioned HDF5 output whose covered CpG loci and methylation ratios are mathematically identical to `methrix` (zero beta discrepancies exceeding 10^*−*6^; Supplementary Note 4, Section S4.4);
- `pairbam, seq2mat, matsrun`, and `qctb` replace the remaining R scripts for mate-pair repair, expression-matrix quantification, rMATS splicing orchestration, and multi-modal QC aggregation, respectively.

## Comparison and practical evaluation

### Functional comparison with existing tools

We compared OTTER with four widely used host-read separation tools (Xenome (Conway et al., 2012), Disambiguate (Ahdesmäki et al., 2016), XenofilteR (Kluin et al., 2018), and Xengsort (Zentgraf and Rahmann, 2021)) across functional and architectural dimensions (Table 1). To our knowledge, OTTER is the only framework that combines bisulfite-aware deconvolution with native strand-aware NM recalculation, an end-to-end multi-omics CLI, zero-dependency static binaries (validated on CentOS 7), asynchronous lock-contention-free Slurm orchestration, and an immutable reference registry.

**Table 1.** Feature and architectural comparison of OTTER with existing host/graft separation tools. Y and N denote presence or absence of each capability.

| Capability / Architectural Dimension | Xenome | Disambiguate | XenofilteR | Xengsort | OTTER (Ours) |
| --- | --- | --- | --- | --- | --- |
| RNA-seq host/graft separation | Y | Y | Y | Y | Y |
| Bisulfite-aware deconvolution (RRBS/WGBS) | N | N | N | N | Y |
| Native strand-aware NM recalculation | N | N | N | N | Y |
| Automated end-to-end multi-omics CLI | N | N | N | N | Y |
| Full-genome CpG HDF5 matrix export | N | N | N | N | Y |
| Zero-dependency static binaries (CentOS 7) | N | N | N | N | Y |
| Asynchronous lock-contention-free Slurm orchestration | N | N | N | N | Y |
| Immutable reference registry & snapshots | N | N | N | N | Y |

### Production deployment and empirical benchmarking

We deployed the release toolchain on two production supercomputing clusters (the Hangzhou Institutional Supercomputing Cluster on CentOS 7/glibc 2.17 and the Paracloud Supercomputing Cluster on Slurm/Lustre storage) and exercised the full pipeline on authentic public SRA cohorts spanning RRBS, RNA-seq, RNA-PDX, and BS-PDX (Supplementary Note 4, Section S4.1). On transcriptomic data, xenofilx achieved exact 100% decision parity with legacy XenofilteR on a 32.1M-fragment RNA-PDX accession and across a 0%–99% RNA mixture gradient (Supplementary Table S7). We therefore focused benchmarking on bisulfite data, comparing `xenofilx` against both the native Picard + `XenofilteR` baseline and the `xdxtools` prototype (Picard `IS_BISULFITE_SEQUENCE=true + XenofilteR`) on ground-truth compositional gradients and an authentic 45.2M-fragment BS-PDX accession (Supplementary Note 4, Section S4.5).

Across a unified 0%–99% BS-seq compositional gradient (43 evaluation cells, Supplementary Table S8), `xenofilx` maintained stable human-graft recall (51.61%–54.03%) at *≈* 99.92% host specificity, consistently outperforming the prototype by +0.63% to +0.70% recall (recovering *≈* 4,000 additional authentic human fragments per million reads in 60% mixtures, and with gains ranging from approximately 7 to 3,914 fragments across the gradient), whereas the naive unpatched baseline collapsed to ***<*** 0.015% recall. Two-stage mathematical decomposition reveals that this apparent 52%–54% recall ceiling is strictly governed by bisulfite-induced C-to-T entropy contraction during primary alignment (*η*_align_ = 54.01%), whereas `xenofilx` achieves 99.23% deconvolution retention on mapped human fragments with ***>*** 99.0% genomic coverage fidelity across 1–10*×* depths (Supplementary Note 4, Section S4.5.4). A 4*×*3 factorial parameter audit (36 runs; Supplementary Table S9) identified mismatch threshold *t* = 7 as the recommended operational bound balancing sensitivity gains against diminishing returns and cross-species alignment noise (52.22% recall, 68.59% F1 at 99.92% specificity), with the unmapped-read penalty invariant across 8– 12. In scheduler evaluations (*n* = 3 paired Slurm repeats per scenario; Supplementary Note 4, Section S4.2), craftmake’s controller reconciliation latency ranged 2.7–4.2 s versus 9.5–19.8 s for the Snakemake compatibility path.

## Discussion and conclusion

OTTER resolves the systemic engineering bottlenecks of the xdxtools prototype (namely dependency fragility, glibc incompatibility, and scheduler lock contention) while preserving its biological foundations, and makes bisulfite-aware xenograft deconvolution practical at production scale. Validation on two production clusters and genuine public SRA datasets spanning RRBS, RNA-seq, RNA-PDX, and BS-PDX establishes OTTER as a reproducible, portable, and high-throughput framework for xenograft multi-omics preprocessing. More broadly, its layered architecture (comprising compiled operators, minimal environments, and deterministic orchestration) offers a general template for hardening script-based bioinformatics pipelines into production systems.

## Acknowledgements

The authors thank Jiamin Sun, Haoyi Wang, and Jian Huang for critical evaluation, testing, and valuable insights during software development.

## Author Contributions

**Yanhua Zheng, Xin Liu**, and **Yingyu Yi**: Conceptualization, Methodology, Software, Validation, Visualization, Writing – Original Draft, Writing – Review & Editing. **Xiaojun Sun** and **Fang Liu**: Data curation, Validation. **Xiaojing Yan, Cheuk-Him Man**, and **Hongcang Gu**: Conceptualization, Investigation, Writing – Review & Editing. **Xiaoxue Wang** and **Fan Zhang**: Funding acquisition, Project administration, Resources, Supervision, Writing – Review & Editing.

## Data Availability

OTTER source code is distributed open-source under the MIT License and is available at https://github.com/otterlab-bio/otter. The legacy prototype is available at https://github.com/Genomiclab/xdxtools. The current reference genome archive is publicly available on Hugging Face (https://huggingface.co/datasets/fallingstar10/xdxtools-genomes). Downsampled SRA test fixtures are provided at https://github.com/otterlab-bio/otter/tree/main/testdata/gate6/craftmake-downsample-20260906/fastq, enabling out-of-the-box reproduction of the evaluated workflows. Test and validation data, including BAM fixtures, mixture truth records, and provenance files, are additionally available at https://huggingface.co/datasets/fallingstar10/otter-data. An agent skill that codifies the recommended OTTER usage and validation workflow (https://github.com/otterlab-bio/otter/tree/main/skills/otter) is also distributed. Public validation and onboarding resources are described in Supplementary Note 4, Section S4.6.

