## Supplementary Information for "OTTER: Orchestrated Transcriptomic, Tumor-xenograft (PDX/CDX), and Epigenomic Reporting Workflow"

#### Supplementary Note 1: The Xdxtools Prototype Architecture and Legacy Workflow

This note documents the architecture, implementation specifications, and operational workflows of the initial proof-of-concept prototype, xdxtools (https://github.com/Genomiclab/xdxtools), which established the biological feasibility of bisulfite-aware deconvolution prior to its systemic overhaul into OTTER.

##### S1.1 Prototype Implementation Overview

The archived xdxtools package (version 0.2.2) declares R ≥ 3.5 and uses an R6 class named BeaverGandalf alongside Snakemake workflow files. The class provides gandalf_create_filework() and gandalf_make_config() methods for workspace/configuration preparation and a gandalf2wars() method for workflow dispatch. During some dispatch paths the object is saved as workflow.RDS, enabling later manual restoration. beaver_engine_detect() identifies the runtime environment (k8s, slurm, or container) by probing command-line binaries in PATH.

##### S1.2 Legacy Input Processing and Phenotype Metadata Contract

The prototype ingested two standardized inputs: paired-end FastQ files formatted as {sampleid}_R1.fastq.gz and {sampleid}_R2.fastq.gz via use_server_fastq(), and experimental metadata sheets via use_server_pdata(). The FastQ ingestion helper listed sequencing files from a designated server path and extracted canonical sample identifiers by stripping read-pair suffixes (suffix1 and suffix2, defaulting to _R1.fastq.gz and _R2.fastq.gz, while allowing user customization in the BeaverGandalf constructor).

Phenotype metadata was provided as a spreadsheet (.xlsx or .xls) loaded via readxl::read_excel(). In xdxtools, the pdata file required specific column structures and naming conventions:

- **sampleid (Required):** A unique sample identifier string that strictly matches the sample prefix resolved from the paired FastQ filenames (with automatic normalization of localized sample identifier headers). We also strictly validate pdata rows against the set of confirmed paired FastQ files (sampleid %in% samples_paired).
- **sample_group (Optional / Recommended):** Experimental cohort, phenotype, or treatment condition annotations (supported aliases include "sample_group", "group", "condition", and "treatment"). BeaverGandalf parsed this column to determine factor levels for downstream alternative splicing analysis; if absent, the class automatically initialized sample_group to NA and issued a single-group notice.
- **inline_barcode_sequence (Optional):** Custom inline barcode sequences for multiplexed sequencing libraries. In BS-seq mode, BeaverGandalf used this column to dynamically generate sample-specific adapter trimming sequences: the reverse complement of the inline barcode was computed and prepended to the 5′ end of standard adapters ("TGA" + revComp + adapter1, and "A" + revComp + adapter2), enabling automated de-multiplexing and adapter trimming in TrimGalore.

After ingesting and sanitizing these inputs, gandalf_create_filework() created the workflow's required filesystem hierarchy, and gandalf_make_config() materialized the parameters into a localized Snakemake YAML configuration.

##### S1.3 Prototype Analytical Pipeline Stages

The archived workflow catalog contains modality-specific Snakemake files for BeaverBS (BS-seq), BeaverPDX (BS-PDX), BeaverRNA (RNA-seq), and BeaverRNASEQPDX (RNA-PDX). Its R scripts orchestrated external bioinformatics utilities across the supported multi-omics sequencing modalities:

- **Step 1 (Quality Control & Trimming):** Raw FastQ streams were evaluated for per-base quality, and GC bias with FastQC + seqkit and trimmed via TrimGalore. Crucially, the prototype enforced a dual-stage quality control protocol: following adapter excision and low-quality base clipping, a second round of FastQC was systematically executed on the post-trimming FastQ files to empirically verify adapter removal efficiency, read length distributions, and quality score restoration prior to sequence alignment.
- **Step 2-1 (Modality-Specific Alignment):** Transcriptomic reads were mapped with STAR (Dobin *et al.*, 2013), while bisulfite reads were mapped with Bismark (Krueger and Andrews, 2011) against composite human and mouse reference genomes.
- **Step 2-2 (Bisulfite-Aware Read Filtration, also designated step2-check):** For xenograft sequencing (BS-PDX, RNA-PDX), we use Picard tools to recalculate alignment tags (NM and NN) and apply C-to-T deamination compatibility via IS_BISULFITE_SEQUENCE=true setting in BS-PDX, followed by XenofilteR (Kluin *et al.*, 2018) disambiguation and automated BAM mate-pair resynchronization (samtools view + repair_unpaired_reads.R + Picard FilterSamReads) to ensure valid paired-end structures.

- **Step 3-1 (Quantification):** Expression counts were extracted using HTSeq (Anders, Pyl and Huber, 2015); methylation calls were parsed into methrix objects (Mayakonda *et al.*, 2020) and exported to bsseq objects (Hansen, Langmead and Irizarry, 2012) via methrix2bsseq() for downstream statistical modeling.
- **Step 3-2 (Reporting, also designated step3-check):** Quality metrics across all pipeline stages (including both rounds of FastQC, TrimGalore, mapping, and disambiguation) were aggregated into a unified report using R scripts (QC_summary.R and QC_summary_rnaseq.R in the xdxtools package).

#### S1.4 Bisulfite Tag Recalibration in the Prototype: Picard IS_BISULFITE_SEQUENCE Audit and Comparative Results

**Tag Recalibration Contract in the Prototype.** In standard bioinformatics pipelines, we use Picard to recalculate alignment-derived edit distance (NM) and mismatch attributes via SetNmMdAndUqTags. In the xdxtools prototype, the deconvolution rule (picard_pdx_patch.smk) explicitly declared:

```
picard SetNmMdAndUqTags \
  I={input.bam_sorted} \
  O={output.fixed_bam} \
  R={params.fasta} \
  IS_BISULFITE_SEQUENCE={params.is_bisulfite}
```

where {params.is_bisulfite} dynamically evaluated to "true" for bisulfite sequencing workflows (BS-seq, RRBS, WGBS) and "false" for transcriptomic workflows (RNASEQ). This parameter configuration was a pivotal algorithmic feature of the prototype: because XenofilteR (Kluin *et al.*, 2018) uses the NM score tag as the primary discriminator for host-
versus-graft disambiguation, the mathematical validity of the deconvolution step depends entirely on whether NM accounts for bisulfite-induced cytosine deamination.

**Empirical Comparison: Enabling vs. Omitting IS_BISULFITE_SEQUENCE.** To quantify the biological and computational impact of this parameter, we systematically benchmarked both configurations across matched bisulfite sequencing datasets (with full production-scale cluster audits compiled in Supplementary Note 4):

1. **Unpatched XenofilteR (IS_BISULFITE_SEQUENCE=false, Unpatched Baseline):** When Picard is executed naively without bisulfite flags against unconverted reference genomes, standard edit-distance algorithms treat every unmethylated cytosine that underwent bisulfite-mediated C-to-T deamination as an alignment mismatch, thus recalculated NM tags inflate to values vastly exceeding the standard mismatch threshold ($t = 6$). As a consequence, XenofilteR rejects almost all authentic graft reads. Across a controlled 60% human BS-seq mixture (1,000,000 fragments, $n = 3$ independent replicates), the unpatched XenofilteR path achieved a human-graft recall of only **0.0068%** (F1 score: 0.0135%; host specificity: 99.9256%), representing a catastrophic sensitivity failure. Across the entire 0% to 99% human content gradient, graft recall remained depressed below 0.015% when deamination discounting was omitted.
2. **Prototype (IS_BISULFITE_SEQUENCE=true, Prototype Path):** Explicitly activating Picard's built-in bisulfite flag discounts valid forward C-to-T deamination events during NM and MD recalculation. On the identical 60% human BS-seq benchmark ($n = 3$ replicates), this restores human-graft recall to **51.3501%** (F1 score: **67.8316%**; host specificity: **99.9184%**), establishing the baseline deconvolution feasibility of the prototype. Across the full 0% to 99% human composition gradient, this prototype configuration achieves stable recall between 50.9067% and 53.3333% with ≈99.92% host specificity (Supplementary Table S8).

| Pipeline Configuration | $t$ | Graft Recall | Host Spec. | Graft PPV | Graft F1 | Retained Frags | Deconvolution Status |
|---|---|---|---|---|---|---|---|
| Unpatched Baseline | 5 | 0.0046% | 99.9258% | 8.4798% | 0.0092% | 325 | Catastrophic collapse |
| Unpatched Baseline | 6 | 0.0068% | 99.9256% | 11.9381% | 0.0135% | 338 | Baseline collapse |
| Unpatched Baseline | 7 | 0.0105% | 99.9253% | 17.3478% | 0.0210% | 362 | Severe under-recovery |
| Unpatched Baseline | 8 | 0.0149% | 99.9249% | 22.9054% | 0.0298% | 390 | Severe under-recovery |
| Prototype | 5 | 50.5244% | 99.9185% | 99.8926% | 67.1069% | 303,472 | Rescued sensitivity |
| Prototype | 6 | 51.3501% | 99.9184% | 99.8942% | 67.8316% | 308,427 | Prototype baseline |
| Prototype | 7 | 51.6993% | 99.9181% | 99.8945% | 68.1357% | 310,524 | Prototype optimal |
| Prototype | 8 | 51.8356% | 99.9177% | 99.8942% | 68.2539% | 311,343 | Prototype plateau |

Table S1: **Impact of Picard IS_BISULFITE_SEQUENCE tag recalibration on prototype bisulfite deconvolution performance.** Evaluated on matched 60% human BS-seq mixtures across three independent 1,000,000-fragment replicates ($n = 3$, containing 600,000 authentic human graft fragments). Unpatched Baseline executes Picard SetNmMdAndUqTags with IS_BISULFITE_SEQUENCE=false; Prototype path enables IS_BISULFITE_SEQUENCE=true as implemented in the xdxtools Snakemake rule. All runs evaluated with unmapped penalty 8. Comprehensive multi-modal benchmarks across the full 0%–99% gradient, modern compiled xenofilx comparisons, and production HPC platforms are documented in Supplementary Note 4 (Supplementary Tables S8 and S9).

#### Supplementary Note 2: OTTER Systems Engineering and Infrastructure Architecture

To systematically overcome the real-world deployment bottlenecks encountered during institutional evaluation of the xdxtools prototype (detailed in Section 2.1), including combinatorial Conda dependency deadlocks, enterprise glibc symbol failures, and shared-filesystem lock contention under parallel execution, OTTER was engineered as a modular, compiled workflow framework. This note details the current product boundaries, one inspected release artifact set on CentOS 7, native DAG orchestration, environment management, and the reference registry.

##### S2.1 Product Hierarchy and Architectural Separation

We structure OTTER into four decoupled compiled layers (a project control plane, an execution layer, an environment lifecycle manager, and specialized domain operators) alongside a foundational BAM/BGZF primitive library shared by the Go operators:

```
otter      Project control plane and canonical-run resolver
  └── craftmake or Snakemake compatibility path
      └── enva-managed environments and workflow operators
          └── bamdriver BAM/BGZF library used by relevant Go operators
```

1. **Control Plane (otter):** The Go CLI coordinates project initialization (otter init), project creation (otter create), canonical run resolution, task inspection, and artifact verification. Canonical snapshots are serialized under the otter.run/v1 schema, while legacy commands continue to support the Snakemake compatibility path. The control plane standardizes and strictly validates two primary user inputs:
    - **Paired-End FastQ Files (--fastq):** Sequencing reads must be supplied as gzip-compressed FastQ files (.fastq.gz or .fq.gz). Paired mates follow the standard naming convention {sampleid}_R1.fastq.gz and {sampleid}_R2.fastq.gz (or custom suffixes specified via --suffix1 and --suffix2). During project creation, the directory scanner verifies mate-pair completeness, confirms non-empty file contents, extracts canonical sample identifiers, and prunes or rejects orphan singleton files.
    - **Phenotype Metadata Table (--pdata):** Experimental and clinical metadata is supplied as an Excel spreadsheet (.xlsx, .xls) or delimited text (.csv). The table requires a sampleid column (with supported aliases including sample_id, sample, and localized variants) whose values must strictly match the parsed FastQ sample strings. For comparative analyses, grouping or treatment conditions are defined via a sample_group or condition column (aliases: sample_group, group, condition, treatment). Multiplexed libraries can optionally include an inline_barcode_sequence column for dynamic adapter calculation. The control plane validates bidirectional consistency between the metadata rows and discovered FastQ pairs, rejecting duplicate identifiers and empty grouping fields prior to snapshot generation.
2. **Execution Layer (craftmake):** The independent Go controller compiles workflow specifications, persists controller state in SQLite, supports local and Slurm backends, and manages task submission, reconciliation, caching, cancellation, and deterministic resumption.
3. **Environment Manager (enva):** The Rust utility uses the rattler ecosystem to create, validate, and invoke named software environments, isolating upstream heavy aligners (Bismark, Bowtie2, STAR) with minimal Conda footprints while providing native takeover and management of environments created by conda, mamba, or micromamba.
4. **Domain Operators:** Statically compiled domain operators (fastqcx, xenofilx, pairbam, seq2mat, matsrun, methx, and qctb) replace the prototype's legacy R tooling across each dedicated computational bottleneck, as detailed in Section 2.3. By packaging read QC, host disambiguation, mate repair, expression matrix quantification, splicing orchestration, methylation calling, and QC aggregation into single-purpose compiled executables, OTTER eliminates R runtime dependencies and pass-by-value memory bloat while preserving validated biological fidelity across a four-stage audit (Gate A: BAM preservation; Gate B: mate integrity; Gate C: NM oracle; Gate D: membership accounting; Supplementary Note 4, Section S4.3).
5. **Foundation Primitives (bamdriver):** The Go library implements high-performance BAM and BGZF reading/writing, coordinate sorting, FASTA-backed bisulfite-aware NM calculation, and alignment-tag handling for downstream Go operators.

##### S2.2 Deployed Release Artifact Verification

On 2026-09-08, we conducted a direct read-only inspection of the accepted release artifacts deployed on the Hangzhou Institutional Supercomputing Cluster host (CentOS Linux 7.6, kernel 3.10.0-957.el7.x86_64, glibc 2.17; compatible across CentOS 7.6–7.9 releases). Executing file and ldd across all eleven installed binaries, we confirmed that none exhibited dynamic runtime dependencies on host shared libraries (Table S2).

| Component | Reported version | Installed size | ldd result on CentOS 7 |
|---|---|---|---|
| otter | daily-20260907+b706fa0 | 13.28 MB | No dynamic dependencies |
| craftmake | 0.1.0-dev+unknown | 8.44 MB | No dynamic dependencies |
| enva | 0.1.0 | 14.85 MB | No dynamic dependencies |
| xenofilx | daily-20260907 | 4.40 MB | No dynamic dependencies |
| pairbam | daily-20260907 | 2.09 MB | No dynamic dependencies |
| seq2mat | daily-20260907 | 5.92 MB | No dynamic dependencies |
| matsrun | daily-20260907 | 4.45 MB | No dynamic dependencies |
| qctb | 0.1.0 | 4.04 MB | No dynamic dependencies |
| fastqcx | 0.3.4 | 24.63 MB | No dynamic dependencies |
| methx | 0.1.0 | 11.29 MB | No dynamic dependencies |
| otter-install | release build | 6.75 MB | No dynamic dependencies |

Table S2: **Verification of the deployed release artifacts on the Hangzhou Institutional Supercomputing Cluster.** Sizes are the installed file sizes observed on 2026-09-08. The ldd column records static linkage verification on the CentOS 7 supercomputing host.

#### S2.3 The Craftmake Execution and Scheduler Engine

craftmake provides a native workflow controller for the documented local and Slurm execution paths. Its persistence, retry, accounting, cache, and compatibility-mode behavior are described below.

**Transactional State Persistence Eliminates Filesystem Lock Protocols.** craftmake stores workflow state in an embedded SQLite database (state.sqlite) and writes controller events to an append-only JSONL log (runs/<run-id>/controller.jsonl). The controller log captures timestamps, run/task/submission/attempt identities, backend job identifiers, statuses, durations, errors, and structured detail fields.

**Controller-First Dispatch Governs Distributed Worker Lifecycles.** When targeting Slurm clusters, craftmake operates under a controller-first topology:

1. **Job Synthesis:** The controller generates self-contained task submission manifests containing explicit CPU core allocations, memory admission limits, time limits, and environment activation wrappers.
2. **Asynchronous Reconciliation:** Task state is queried through periodic Slurm accounting and queue queries (sacct and squeue). On worker completion, exit status and available resource metrics are persisted in state.sqlite.
3. **Graceful Signal Handling:** On SIGTERM or SIGINT, the controller requests cancellation of outstanding Slurm allocations through scancel, records the resulting state transition, and preserves controller evidence.

#### S2.4 Lightweight Environment Management with Enva

**Conda-Compatible Environment Isolation without External Package Manager Overhead.** While domain operators ship as self-contained static binaries, upstream aligners (Bismark, Bowtie2, STAR) require a resolved scientific software environment. We build enva to manage software environments partitioned by operational necessity without requiring an external conda, mamba, or micromamba installation:

- **otter-core (Production Runtime):** The sole required runtime environment for standard OTTER production workflows, encapsulating essential upstream aligners and core utilities (Bismark 3.1.0, Bowtie2 2.5.5, Samtools 1.24, STAR 2.7.11b, and rmats.py 4.4.0) with a minimized dependency footprint.
- **otter-snakemake (Regression Testing):** Maintained specifically for parity verification and regression testing of legacy Snakemake workflows against native craftmake execution.
- **otter-extra (Legacy Compatibility):** Reserved for historical compatibility audits with the legacy R package collection inherited from the prototype.

#### S2.5 Centralized, Immutable Reference Genome Registry

**Cryptographic Manifests and Atomic Publication Seal Genomic Assets.** In legacy pipelines, the lack of centralized reference genome management forced users to manually specify paths to FASTA and GTF files. This manual approach became particularly chaotic and error-prone when handling multiple species across diverse assembly and annotation versions. OTTER introduces an immutable Reference Genome Registry ($OTTER_REFERENCE_ROOT):

```
$OTTER_REFERENCE_ROOT/
└── genomes/<reference-id>/<release>/
    ├── reference.yaml       # Formal metadata specification (otter.reference/v1)
    ├── manifest.json        # Cryptographic manifest of all constituent files
    ├── checksums.sha256     # Line-delimited SHA-256 validation manifest
    ├── fasta/
    │   ├── <reference-id>.fa
    │   └── <reference-id>.fa.fai
    ├── annotations/
    │   └── <reference-id>.gtf
    └── indexes/
        ├── bismark/         # Bismark CT/GA converted reference indexes
        ├── bowtie2/         # Bowtie2 index files
        └── star/            # STAR index files
```

Analysis projects bind to references via references.lock.yaml, recording only logical reference identifiers and their manifest SHA-256 digest:

```yaml
schema_version: otter.references.lock/v1
references:
  primary:
    id: hg19
    release: hg19-legacy-ensGene
    manifest_digest: sha256:f6021930ac2017803e612f57f32347337b32ad016b9570f0a43eb750a730fc98
```

To facilitate migration from legacy deployments, we provide scripts/migrate_legacy_reference.sh in OTTER. In --mode ingest, the utility hardlinks existing legacy Bismark and STAR indexes, builds samtools FAI indexes, generates verified reference.yaml / manifest.json manifests, and seals the release in under 2 minutes without re-indexing.

#### S2.6 Self-Contained Go Installer Distribution (otter-install)

In deployment, we provide otter-install, a single-binary installer implemented as a Go module. Upon execution, otter-install:

1. Queries release metadata and downloads the ten managed operator binaries to ~/.cargo/bin;
2. Automatically unpacks and deploys the embedded workflow catalog to ~/.cargo/share/craftmake/workflows/ and configuration templates to ~/.cargo/share/craftmake/configs/;
3. Invokes each managed tool's --version contract where available; and
4. Supports an optional GitHub proxy prefix (--github-proxy) for firewalled institutional networks.

#### Supplementary Note 3: Mathematical Modeling and Algorithmic Formulations

To ensure reproducible systems engineering and mathematical transparency, this note details the strand-aware bisulfite NM and classification rules in xenofilx, the current chromosome-bucket annotation procedure in methx, and the versioned custom HDF5 storage layout (methx.custom-hdf5/1.0.0).

##### S3.1 Reference-Aware Bisulfite NM and Classification Formulation

**Chemical Cytosine Deamination Induces Strand-Asymmetrical Transitions.** Sodium bisulfite treatment chemically deaminates unmethylated cytosines into uracils, which are subsequently amplified as thymines during polymerase chain reaction (PCR). Converted reads therefore contain directional base transitions that must not be scored as ordinary mismatches.

Let $R = (r_1, r_2, ..., r_L)$ be the reference sequence over the aligned window, $S = (s_1, s_2, ..., s_K)$ be the read sequence, and $\mathcal{C}$ be its SAM CIGAR operations (Li *et al.*, 2009). The current bamdriver implementation first checks the Bismark XG tag. For a valid XG=CT record, only $\mathrm{C} \to \mathrm{T}$ is exempted; for a valid XG=GA record, only $\mathrm{G} \to \mathrm{A}$ is exempted. If this tag is unavailable, the fallback orientation indicator $\sigma$ is derived from SAM FLAG 0x10:

$$\sigma = \begin{cases} 0 \text{ if } (\mathrm{FLAG} \wedge 0\mathrm{x}10) = 0 & (\text{forward fallback}) \\ 1 \text{ if } (\mathrm{FLAG} \wedge 0\mathrm{x}10) = 0\mathrm{x}10 & (\text{reverse fallback}) \end{cases}$$

The fallback conversion indicator is:

$$\mathbb{I}_{\mathrm{bs}}(r_i, s_j, \sigma) = \begin{cases} 1 \text{ if } \sigma = 0 \wedge r_i = \mathrm{C} \wedge s_j = \mathrm{T} \\ 1 \text{ if } \sigma = 1 \wedge r_i = \mathrm{G} \wedge s_j = \mathrm{A} \\ 0 \text{ otherwise} \end{cases}$$

**Basewise CIGAR Inspection Derives Strand-Conditioned Edit Distance.** For CIGAR M operations, the implementation counts non-identical aligned bases unless the appropriate conversion indicator holds. For CIGAR X, conventional mode counts the operation length; bisulfite mode first counts that length and then subtracts basewise conversion-compatible positions. Insertions and deletions each contribute their CIGAR length, yielding:

$$\mathrm{NM}_{\mathrm{bs}} = (N_{\mathrm{mis}} - N_{\mathrm{exempt}}) + \sum_{o \in \mathcal{C}_{\mathrm{ins}}} \mathrm{len}(o) + \sum_{o \in \mathcal{C}_{\mathrm{del}}} \mathrm{len}(o)$$

where $N_{\mathrm{mis}}$ represents the total count of raw mismatched base pairs across aligned segments (under M and X operations), and $N_{\mathrm{exempt}}$ denotes the subset of conversion-compatible transitions matching indicator $\mathbb{I}_{\mathrm{bs}}(r_i, s_j, \sigma)$.

**Inherited Score Heuristics Retain Upstream Classification Compatibility.** xenofilx retains the inherited XenofilteR scoring convention:

$$S_{\mathrm{alignment}} = \mathrm{NM}_{\mathrm{bs}} + N_{\mathrm{ins}} + N_{\text{soft clip}}$$

Thus, insertion bases are represented both within the SAM-style NM term and as a separate inherited classification-score component. This explicit compatibility formulation deliberately preserves XenofilteR's compound ranking metric rather than enforcing a strict edit-distance interpretation.

**Conditional Disambiguation Gating Enforces Mate-Level Quality Bounds.** For a paired fragment, each mate must meet the configured graft mismatch threshold before a graft assignment can be considered. When both species have usable primary pairs, the classifier compares the sums of the two mate scores and assigns the lower total; equal totals are discarded. When the host pair is absent, graft assignment still requires that at least one graft mate score is below the configured threshold. This dual-mate gating protocol guarantees fragment-level disambiguation fidelity, providing the algorithmic foundation for robust host disambiguation prior to downstream genomic feature annotation.

##### S3.2 Pre-Compiled CpG Extraction and Chromosome-Bucket Annotation in Methx

**Pre-Compiled CpG Coordinate Extraction into Serialized RON Format.** In legacy R workflows, extracting reference CpG coordinates via methrix::extract_CPGs() required repetitive runtime parsing of multi-gigabyte genomic FASTA files or heavy BSgenome packages, consuming extensive execution time and multi-gigabyte memory on every cohort run. methx replaces this bottleneck with an ahead-of-time pre-compilation step (methx extract-cpgs -g <genome.fa> -o <reference_cpgs.ron>). The Rust extractor streams genomic FASTA files (supporting uncompressed .fa and multi-member gzipped .fa.gz streams), parses canonical contigs without loading the entire genome into memory, identifies all forward and reverse CpG dinucleotides, and records their exact 0-based coordinate intervals and strand orientation (CpGSite { chr, start, end, strand }) alongside contig lengths.

The resulting CpGData structure is serialized into Rusty Object Notation (.ron) and written atomically to disk (reference_cpgs.ron). In downstream analysis (methx process), the pipeline deserializes this pre-compiled coordinate registry in seconds, eliminating repetitive genome extraction across pipeline invocations.

**Chromosome-Specific 64-KiB Bucketing Accelerates Localized Feature Traversal.** For genomic feature annotation, methx parses reference GTF annotations into transcript-derived genomic intervals (promoters, 5′/3′ UTRs, exons, introns, downstream regions, and intergenic spaces). To accelerate spatial overlap queries during methylation quantification, it partitions feature intervals into chromosome-specific 64-KiB genomic buckets. For any queried CpG position $p$, the engine derives the spatial bucket index $b = \lfloor \frac{p}{2^{16}} \rfloor$, retrieves the candidate intervals within bucket $b$, and determines containment:

$$\text{contains}([s_i, e_i), p) \text{ iff } s_i \le p < e_i$$

When multiple annotations overlap, a deterministic priority cascade is enforced:

$$\text{promoter} > \text{UTR} > \text{exon} > \text{intron} > \text{downstream} > \text{intergenic}$$

Ties within the same priority class are resolved by selecting the closest transcription start site (TSS). For non-overlapping coordinates, a binary partition over ordered TSS positions combined with a localized sliding window assigns the nearest-gene metadata. Feature traversal is parallelized across available threads via Rayon, producing comprehensive CpG coverage statistics (CpG_coverage.xlsx) and split genomic region reports without runtime R interpreter overhead.

##### S3.3 Versioned Custom HDF5 Storage Contract

**Bounded Chunk Serialization Precludes Full-Matrix In-Memory Transposition.** The current methx.custom-hdf5/1.0.0 writer produces an atomic custom HDF5 file containing the following datasets:

```
/
├── beta                  # Float32 assay: shape (samples, CpGs)
├── cov                   # UInt32 assay: shape (samples, CpGs)
├── rowData/
│   ├── chr, seqnames          # Variable-length chromosome strings
│   ├── start, end, width      # UInt32, 1-based closed coordinates
│   └── strand               # Variable-length '+', '-', or '*' strings
├── colData/
│   ├── sample_id
│   └── sample_name
└── metadata/
    ├── genome
    ├── schema_name            # "methx.custom-hdf5"
    ├── schema_version         # "1.0.0"
    ├── loader_compatibility
    └── is_h5
```

Assays are written one sample/CpG block at a time, with shape $(S, M)$, chunk shape $(1, \min(262144, M))$, and deflate compression level 6. This bounded writer avoids a complete in-memory transpose before publication. The staged file is validated by the Rust-native schema/readback validator before atomic publication.

**Direct Dataset Queryability and Downstream Methrix Reconstitution.** The primary storage contract provides direct rhdf5 access to the versioned datasets (/beta, /cov, /rowData, /colData, /metadata). To preserve atomic single-file distribution on shared filesystems, the output is serialized as a self-contained .h5 archive rather than a multi-file saveHDF5SummarizedExperiment() directory bundle from the methrix package. To ensure seamless interoperability with the Bioconductor methylomics ecosystem, we provide a dedicated R loader function (load_h5_as_methrix) that parses the versioned assays, constructs a genomic GRanges coordinate index from rowData, binds sample annotations from colData, and reconstitutes a fully compliant SummarizedExperiment and methrix object. This verified loader function confirms that methx output can be directly loaded into downstream R analytical workflows (including methrix and bsseq) without intermediate text formatting or coordinate realignment.

### Supplementary Note 4: Comprehensive Empirical Benchmarks and Production Validation

This note compiles the empirical benchmarks, operational cluster verifications, and algorithmic quality audits obtained across our two production high-performance computing (HPC) environments and authentic public multi-omics sequencing cohorts. Section S4.6 separately describes public software and data distribution, continuous integration, and interactive onboarding, distinguishing engineering contract checks from the scientific evaluations reported in Sections S4.1–S4.5.

#### S4.1 Production High-Performance Computing (HPC) Infrastructure and Public SRA Benchmark Corpus

To evaluate OTTER under real-world supercomputing conditions, we deployed and benchmarked the framework across two distinct production HPC environments:

1. **Hangzhou Institutional Supercomputing Cluster:** An enterprise legacy Linux cluster operating on CentOS Linux 7.6 (x86_64) with Linux kernel 3.10.0-957.el7.x86_64 and GNU C Library glibc 2.17 (compatible across CentOS 7.6–7.9). The Hangzhou cluster served as the baseline target for verifying zero-dynamic-dependency static linkage(ldd audit across all 11 release binaries, Supplementary Table S2),reference genome migration, and single-node compute execution.
2. **Paracloud Supercomputing Cluster:** A high-throughput cloud supercomputing facility operating on CentOS Linux release 7.9.2009 Core (x86_64) with Linux kernel 3.10.0-1160.118.1.el7.x86_64, glibc 2.17, and Slurm 23.11.8. Login nodes feature Intel Xeon CPU E5-2620 v4 processors (@ 2.10 GHz) interconnected via high-speed InfiniBand networking and backed by a high-performance Lustre parallel shared filesystem mounted across cluster compute nodes. Paracloud served as the primary testbed for distributed scheduler benchmarking, full-pipeline multi-omics execution, and host-read deconvolution testing.

**Enterprise Linux Deployment Verification on the Hangzhou Supercomputing Host.** To verify real-world cluster portability on legacy enterprise Linux environments, we performed a read-only deployment inspection on the Hangzhou Institutional Supercomputing Cluster host on 2026-09-08. Our ldd audit confirmed zero dynamic runtime dependencies across all eleven inspected deployed artifacts (the ten operator binaries plus otter-install, Supplementary Table S2). Querying the deployed otter-core environment, we confirmed Bismark 3.1.0, Bowtie2 2.5.5, Samtools 1.24, STAR 2.7.11b, and rmats.py 4.4.0.

**Operational Traceability across Ingestion, Dispatch, and Cache Resumption.** Our accepted operational log records: (1) we ingested an existing hg19 reference bundle (3.0 GB FASTA, 448 MB GTF, with Bismark, Bowtie2, and STAR assets) using scripts/migrate_legacy_reference.sh --mode ingest, completing manifest validation and read-only sealing without redundant disk duplication; (2) we executed a minimal BeaverBS/step1 workflow dispatching fastqc_before, trim_reads, and fastqc_after to compute node comput3 within partition cpu112c; and (3) we performed a subsequent craftmake resume that registered three cache hits (decision=hit).

**Public SRA Multi-Omics Workflow Validation Corpus.** To validate the end-to-end OTTER workflows across real-world sequencing scenarios, we utilized authentic public Sequence Read Archive (SRA) datasets spanning all four supported multi-omics modalities: reduced representation bisulfite sequencing (RRBS), transcriptomic RNA sequencing (RNA-seq), RNA patient-derived xenograft (RNA-PDX), and bisulfite patient-derived xenograft (BS-PDX; Table S3). Across these seven accessions, the workflow executed complete processing pipelines and generated nine canonical alignment-BAM fixtures to benchmark pipeline execution, toolchain parity, and disambiguation fidelity:

- **Human Reduced Representation Bisulfite Sequencing (RRBS, SRR31480456):** Primary human bisulfite sequencing, aligned against hg19 / hg38 (1 canonical BAM); exercised through end-to-end step1 to step3-check, validating Bismark mapping, methx custom HDF5 CpG matrix generation, and qctb quality aggregation.
- **Mouse Reduced Representation Bisulfite Sequencing (RRBS, SRR10025242):** Murine bisulfite sequencing, aligned against mm10 (1 canonical BAM); provides a single-species bisulfite validation baseline.
- **Human Transcriptomic RNA-seq (SRR1039508):** Paired-end mRNA-seq of human airway smooth muscle cells (HASMC), aligned against hg38 (1 canonical BAM); utilized for conventional read-level NM oracle blind auditing, a 50:50 synthetic RNA mixture pilot, and compositional gradient benchmarks.

- **Human Transcriptomic RNA-seq (SRR018258):** Paired-end human transcriptomic sequencing, aligned against hg38 (1 canonical BAM); exercised through full quantification, MultiQC aggregation, and Slurm walltime incident recovery testing.
- **Mouse Transcriptomic RNA-seq (SRR037954):** Paired-end murine transcriptomic sequencing, aligned against mm10 (1 canonical BAM); paired in silico with SRR1039508 to generate synthetic chimeric sequencing mixtures.
- **Patient-Derived Xenograft Transcriptomic RNA-seq (RNA-PDX, SRR30880970):** Paired-end sequencing of human tumor graft tissue infiltrating murine host stroma; simultaneously aligned against hg38 (human graft) and mm10 (mouse host) to produce 2 canonical alignment-BAM fixtures; evaluated for transcriptomic host-read deconvolution (xenofilx) and Slurm controller reconciliation latency ($n = 3$ paired repeats).
- **Patient-Derived Xenograft Bisulfite Sequencing (BS-PDX, SRR36187610):** Authentic high-throughput enhanced RRBS (eRRBS) sequencing of human tumor xenograft tissue containing 45,160,030 fragments and 10,033,858 covered CpG loci; simultaneously aligned against bisulfite-converted hg38 and mm10 to produce 2 canonical alignment-BAM fixtures; evaluated for bisulfite-aware host read disambiguation (xenofilx --bisulfite), Gate D fragment classification accounting, full-genome methylation matrix extraction (methx), and Slurm controller reconciliation latency ($n = 3$ paired repeats).

Following benchmark standardization, SRR36187610 serves as the definitive BS-PDX benchmark corpus providing complete genome-wide eRRBS coverage, while downsampled fixtures from preliminary accession SRR23802966 are retained solely for local unit-test execution.

| Accession | Modality | Organism / Source | Target References | Primary Analytical Role in OTTER |
|---|---|---|---|---|
| SRR31480456 | Human RRBS | **H. sapiens** (primary tissue) | hg19 / hg38 | End-to-end bisulfite processing and human source for the 0%–99% BS-seq gradient |
| SRR10025242 | Mouse RRBS | **M. musculus** (tissue) | mm10 | Murine bisulfite baseline and mouse source for the 0%–99% BS-seq gradient |
| SRR1039508 | Human RNA-seq | **H. sapiens** (airway smooth muscle, HASMC) | hg38 | Conventional NM oracle blind audit and assay-specific RNA mixture control |
| SRR018258 | Human RNA-seq | **H. sapiens** (transcriptome) | hg38 | MultiQC reporting and transient Slurm walltime recovery testing |
| SRR037954 | Mouse RNA-seq | **M. musculus** (brain tissue) | mm10 | Paired with SRR1039508 for the separate conventional RNA mixture control |
| SRR30880970 | RNA-PDX | Human tumor graft in mouse host | hg38 (graft) + mm10 (host) | Dual-species STAR alignment (2 BAMs), deconvolution, Slurm scheduler repeats |
| SRR36187610 | BS-PDX (eRRBS) | Human tumor graft in mouse host | hg38 (graft) + mm10 (host) | Dual Bismark alignment (2 BAMs), Gate D accounting, full CpG matrix extraction |

Table S3: **Characteristics and analytical roles of the public SRA benchmark corpus.** Seven genuine public SRA accessions represent the four core multi-omics scenarios (RRBS, RNA-seq, RNA-PDX, and BS-PDX). In xenograft scenarios, each library is aligned against both human graft and mouse host reference genomes, yielding nine canonical alignment-BAM fixtures.

##### S4.2 Craftmake vs. Snakemake: Controller Reconciliation Latency

**Controller Reconciliation Latency Isolates Orchestrator Coordination Overhead.** To evaluate whether craftmake mitigates the filesystem lock contention documented in Snakemake (Section 2.1), we benchmarked the **controller reconciliation latency** of the step2-check workflow phase under real Slurm cluster workloads on the Paracloud Supercomputing Cluster. Controller reconciliation latency measures the exact wall-clock interval from the physical completion of the final worker job on a compute node to the controller's recorded terminal DAG completion. Given the execution demands of full-scale Slurm workloads, controller reconciliation latency was evaluated across $n = 3$ paired repeats to characterize median overhead and empirical dispersion (exact two-sided Wilcoxon signed-rank $p = 0.250$, the minimum achievable value at $n = 3$).

**Balanced Cluster Executions Across Multi-Omics PDX Scenarios.** The experiment evaluated six balanced executor pairs: three paired repeats for reduced-representation bisulfite PDX (bs-pdx) and three paired repeats for RNA-seq PDX (rna-pdx) (Table S4 and Figure S1). Each pair evaluated identical inputs, reference manifests, toolchain binaries, and Slurm resource envelopes.

| Scenario | Repeats | Craftmake Median (Min–Max), s | Snakemake Median (Min–Max), s | Paired Diff, s | Wilcoxon $p$ |
|---|---|---|---|---|---|
| BS-PDX | 3 | 4.213 (2.385–5.057) | 19.819 (14.008–164.668) | 15.606 | 0.250 |
| RNA-PDX | 3 | 2.651 (1.552–4.308) | 9.478 (8.458–29.556) | 5.807 | 0.250 |

Table S4: **Controller reconciliation latency comparison between Craftmake and Snakemake.** Evaluated across balanced Slurm runs on identical cluster hardware on the Paracloud Supercomputing Cluster. Paired difference represents the median paired difference (Snakemake − Craftmake). Exact two-sided Wilcoxon signed-rank $p = 0.250$ represents the minimum achievable $p$-value for $n = 3$ paired repeats.

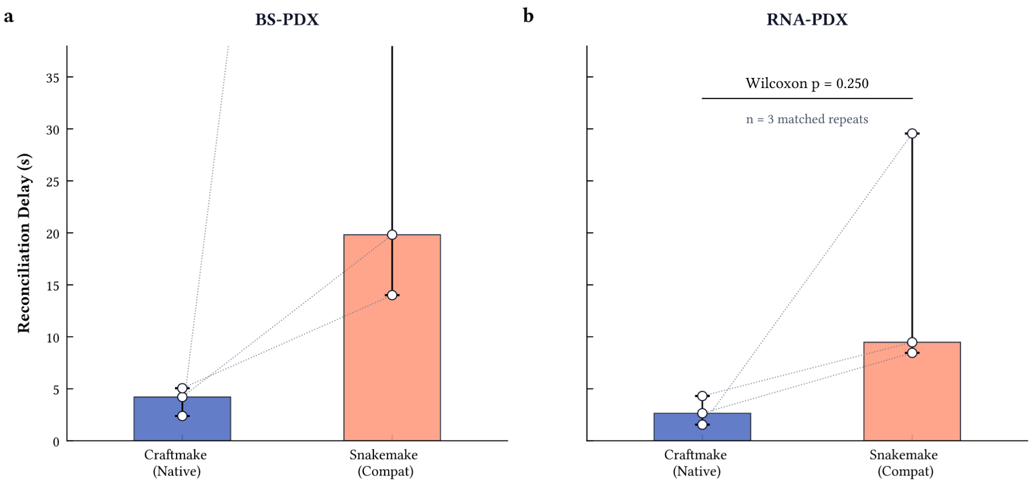

Figure S1: **Median controller reconciliation latency and observed ranges across paired Slurm repeats.** Bars depict the median reconciliation duration across three paired executions on the Paracloud Supercomputing Cluster. Whiskers indicate the empirical minimum and maximum ranges, and connecting gray lines link matching repeats.

**Empirical Tail-Latency Containment Without Distributed Lock Contention.** Across these paired Slurm evaluations, Craftmake reconciliation latency exhibited medians of 4.213 s (BS-PDX) and 2.651 s (RNA-PDX), compared with 19.819 s and 9.478 s for the Snakemake compatibility projection. The observed Snakemake maximum was 164.668 s and the Craftmake maximum was 5.057 s.

##### S4.3 Bamdriver-Based Processing and Quality Audits: Gate A–D Matrix and Fastqcx Parity

The OTTER operator suite was validated through a four-stage quality audit protocol (Gates A through D) using authentic sequencing datasets on the Paracloud Supercomputing Cluster (Table S5).

| Gate | Subject | Evaluation Scope | Audit Criteria | Outcome |
|---|---|---|---|---|
| Gate A | BAM Preservation (bamdriver) | 90,320,060 graft + 1,885,462 host real BAM records | Lossless decode/encode; canonical fields, headers, tags; samtools quickcheck | **100% Pass** |
| Gate B | Mate Integrity (pairbam) | 942,731 primary fragments (1,885,462 records) | Canonical mate resolution, pointer/TLEN preservation, selection subsetting | **100% Pass** |
| Gate C | NM Oracle Blind Test (xenofilx) | 200,000 read prefix per cell across RNA, BS-PDX host, and BS-PDX graft | Independent oracle traversal vs Xenofilx score; converted-reference controls | **100% Pass** (200k/200k per cell) |
| Gate D | PDX Decision Accounting | 45,160,030 fragments from genuine PDX eRRBS cohort (SRR36187610) | 44,044,790 agreements; complete accounting of 1,115,240 membership differences | **Closed with limitation** |

Table S5: **Summary of the four-stage Gate A–D systems validation matrix.** Gates A–C validate the declared preservation and score-audit contracts. Gate D closes a complete accounting of observed membership differences under legacy/modern semantic limitation, and we perform a detailed analysis in Supplementary Note 4, Section S4.5.

**Gate A (BAM Decode/Encode Preservation):** We validated bit-level BAM preservation on authentic production BAM files (39 m 30 s wall-clock). Decoded headers, coordinate sequences, base qualities, and auxiliary tags matched the input streams with zero record losses or unexpected mutations across all 92,205,522 records (90,320,060 graft and 1,885,462 host). We verified that both round-trip BAM outputs passed samtools quickcheck.

**Gate B (Pairbam Preservation and Mate Integrity):** We evaluated mate integrity on the frozen mm10-mapped host BAM (4 m 43 s wall-clock). We confirmed that pairbam retained all 942,731 complete primary fragments (1,885,462 records) with 100% mate-pointer and template-length fidelity.

**Gate C (Read-Level NM Oracle Blind Test):** We performed a blind oracle audit using an independently coded CIGAR/SEQ/reference validation oracle sharing no implementation code with xenofilx or bamdriver. Across 200,000-record prefixes, we observed:

1. Conventional RNA (SRR1039508): Xenofilx non-bisulfite NM and Picard conventional NM matched our independent oracle across all 200,000/200,000 records.
2. BS-PDX host mm10: Xenofilx strand-aware bisulfite NM matched the corrected oracle across all 200,000/200,000 records. Converted-reference control arms (XG=CT and XG=GA) each matched the converted-reference oracle across 200,000/200,000 records.
3. BS-PDX graft hg38: Xenofilx strand-aware bisulfite NM matched the corrected oracle across all 200,000/200,000 records, with matching control arms.

**Gate D (PDX Membership Difference Accounting):** Gate D was evaluated as a fragment-level membership comparison, not as a comparison of BAM record counts. Both pipelines received the same frozen paired-species BAM inputs and the same fragment universe. The modern path was xenofilx with --recalculate-nm --bisulfite, native strand-aware NM recalculation through bamdriver, --mm-threshold 6, --unmapped-penalty 8, and an 8G sort-memory budget in the accepted direct comparison run. The prototype-equivalent path split alignments by strand, recalculated tags with Picard SetNmMdAndUqTags against C-to-T or G-to-A converted references, merged the strand partitions, and ran the original R XenofilteR implementation with the same mismatch threshold (6) and unmapped-read penalty (8). Thus, the comparison changes the classifier implementation and NM/tagging path while holding the input, references, and decision parameters fixed.

For the genuine BS-PDX eRRBS cohort SRR36187610 (graft hg38, host mm10), the direct comparison covered 45,160,030 fragments. The modern and prototype paths agreed on 44,005,684 graft/graft fragments and 39,106 discarded/discarded fragments. They differed for 1,115,240 fragments: 1,115,239 modern-only graft calls (retained as human graft by xenofilx, discarded by Picard + XenofilteR) and one prototype-only graft call (retained by Picard + XenofilteR, discarded by xenofilx). The difference is therefore a bounded membership-semantic difference, not a missing-input or BAM-corruption result.

Dedicated fragment reconciliation partitioned every modern-only call: 1,025,732 had no murine host primary pair and passed the modern graft threshold; 88,923 had complete graft and host primary pairs and a lower modern graft composite score than the host score; and 584 were discarded under both paths for different internal reasons. These categories sum exactly to 1,115,239, with zero missing score records and zero unexplained disagreements. Gate D is consequently closed with prototype/modern semantic limitation.

**Real-world sequencing scenarios comparison: PDX assay-specific parameter contracts.** BS-PDX and RNA-PDX were not evaluated with interchangeable settings. RNA-PDX SRR30880970 used the production RNA threshold 4, penalty 8, ordinary-reference conventional NM, and omitted --bisulfite, achieving 100% fragment-level agreement (32,097,614/32,097,614 fragments) between xenofilx and Picard + XenofilteR. In contrast, BS-PDX SRR36187610 used threshold 6, penalty 8, and strand-aware bisulfite NM, including --bisulfite in xenofilx and strand-partitioned CT/GA Picard recalculation in the prototype-equivalent path, with all 1,115,240 observed differences fully accounted for.

| Assay | Inputs and parameters | Modern path | Prototype path | Membership |
|---|---|---|---|---|
| RNA-PDX | SRR30880970; hg38 graft + mm10 host; threshold 4; unmapped penalty 8 | xenofilx; --recalculate-nm; conventional NM; no --bisulfite | Picard SetNmMdAndUqTags on references fasta + R XenofilteR | 32,097,614 fragments; 32,097,614 agreements; 0 differences; 30,773,947 graft/graft and 1,323,667 discarded/discarded |
| BS-PDX | SRR36187610; hg38 graft + mm10 host; threshold 6; unmapped penalty 8 | xenofilx; --recalculate-nm --bisulfite; strand-aware NM | Strand-partitioned Picard SetNmMdAndUqTags on references fasta + R XenofilteR | 45,160,030 fragments; 44,044,790 agreements; 1,115,240 differences |

Table S6: **PDX membership-comparison evidence and assay-specific parameter contracts.** The RNA-PDX cell uses the lower production RNA threshold and ordinary-reference conventional NM, establishing complete fragment agreement. The BS-PDX cell records the accepted strand-aware bisulfite comparison on the identical frozen fragment universe with prototype/modern semantic limitation.

**Quality Control Parity between fastqcx, FastQC, and SeqKit.** Beyond BAM and deconvolution operations, we verified upstream read quality control parity on identical raw and trimmed FastQ inputs across the benchmark corpus against the legacy oracle combination (FastQC 0.12.1 and SeqKit 2.13.0). Modern compiled fastqcx produced exact numerical agreement across all 16 SeqKit summary fields (total read counts, sequence length bounds, overall GC percentages, and Q20/Q30 fractions). On FastQC-specific analytical modules, evaluation across authentic RNA-PDX SRR30880970 fixtures and synthetic repetition controls demonstrated exact numerical agreement across all 55 per-base quality position groups (330 values, maximum difference 0.0), all 16 sequence duplication bins (deduplicated percentage 94.97%, maximum difference 0.0), and overrepresented sequence identities and abundances. Modules outside this audited scope (such as adapter content and per-tile quality distributions) remain deferred to downstream inspection. This confirms that fastqcx consolidates multi-tool QC metrics into a single high-performance pass with bounded, contract-verified parity.

#### S4.4 Methx: Empirical Annotation Throughput, Memory Benchmarking, and Methrix Parity

**CpG Universe and Production Matrix Parity between Methx and Methrix.** We audited methx against the reference methrix Bioconductor package on identical frozen inputs, utilizing the public SRR36187610 eRRBS dataset and the hg38 reference assembly. Across both implementations, the reference CpG catalog comprised 29,401,795 loci across 25 canonical contigs. Under standard production parameters (min_coverage = 1, remove_uncovered = true, and four execution threads in methx; stranded = TRUE, collapse_strands = TRUE, pipeline = "Bismark_cov", and zero_based = FALSE in methrix 1.8.1), both tools filtered 19,367,937 zero-coverage loci and retained an identical universe of 10,033,858 covered CpG rows. Across all 10,033,858 genomic loci, coordinate sets and sample identifiers matched with 100% exact parity, coverage value mismatches were zero, beta methylation discrepancies exceeding $10^{-6}$ were zero (maximum absolute beta difference was $2.97 \times 10^{-8}$ attributable to single-precision float32 arithmetic), and matrix cell discrepancies were zero. Native Bioconductor interoperability was further established by exporting the methx HDF5 output and seamlessly reloading it with methrix::load_HDF5_methrix().

**Comparative Profiling: Canonical End-to-End Workflow vs. Pre-Filtered Reference Extraction.** In standard bioinformatics practice, researchers execute methrix::extract_CPGs(ref_genome = "BSgenome.Hsapiens.UCSC.hg38") to dynamically instantiate the whole-genome CpG universe prior to populating methylation matrices. To evaluate authentic production performance under identical cluster allocations (four dedicated CPU cores, 128 GiB memory ceiling), we benchmarked the complete canonical workflow of both engines on the SRR36187610 cohort. Under this canonical contract, methx completed in 1 minute 32 seconds (91.6 s) with a peak RSS of 6.42 GiB, extracting reference CpGs directly from FASTA and streaming the versioned HDF5 matrix to disk. In contrast, the canonical R Methrix workflow required 3 minutes 41 seconds (221.0 s) and expanded to 18.62 GiB peak RSS. Across pipeline phases, Methrix expended 56.2 seconds extracting 29,401,795 reference CpGs from the BSgenome package, 131.2 seconds in read_bedgraphs (aligning 58,803,590 stranded positions), 3.2 seconds in remove_uncovered, and 19.8 seconds in HDF5 serialization. Profiling reference extraction in isolation revealed that merely instantiating the reference CpG coordinates in R consumes 4.14 GiB peak RSS and 55.6 seconds before processing any sequencing reads. Against this canonical standard, methx achieves a 2.41-fold wall-clock speedup (91.6 s vs. 221.0 s) and reduces peak memory consumption by 65.51% (6.42 GiB vs. 18.62 GiB).

By comparison, in a restricted covered-matrix evaluation where Methrix was supplied with pre-extracted coordinates (bypassing runtime whole-genome reference extraction), memory-resident execution in R completed in 1 minute 3 seconds at 5.65 GiB peak RSS. Under this narrow scope, methx recorded 6.42 GiB peak RSS (13.75% higher) because it concurrently performs whole-genome reference extraction on the fly rather than requiring pre-computed coordinate files. Post-run matrix auditing across all 10,033,858 covered rows confirmed 100% numerical concordance across all evaluated workflows.

#### S4.5 Xenofilx: Multi-Omics Host Deconvolution Parity, Epigenomic Sensitivity, and Parameter Response

**Methodological Separation and Assay-Specific Contracts.** RNA-seq and bisulfite sequencing cannot be evaluated under interchangeable deconvolution parameters or aggregated into a single composite performance curve. Transcriptomic host deconvolution relies on spliced STAR alignments evaluated under ordinary-reference edit distance, where standard NM metrics apply directly. Conversely, bisulfite sequencing operates on chemically deaminated reads mapped via Bismark to bisulfite-converted genomes, where unmethylated cytosines are converted to uracils (thymines). Because this asymmetric deamination reduces sequence entropy, standard reference-dependent edit distance misinterprets valid conversion events as mismatches, fundamentally altering alignment score distributions. Consequently, we establish two distinct production contracts: RNA-PDX uses --mm-threshold 4, --unmapped-penalty 8, and conventional NM recalculation (--recalculate-nm without --bisulfite), whereas BS-PDX uses --mm-threshold 6, --unmapped-penalty 8, and native strand-aware bisulfite NM (--recalculate-nm --bisulfite). To rigorously characterize deconvolution behavior across both domains, we first evaluate baseline algorithmic parity in transcriptomics (authentic RNA-PDX and a 0%–99% RNA mixture gradient), proceed to bisulfite epigenomic deconvolution (authentic BS-PDX and a 0%–99% BS-seq gradient), and finally conduct a systematic factorial parameter sensitivity analysis on bisulfite deconvolution.

##### *S4.5.1 RNA-PDX and Transcriptomic Deconvolution Parity across 0% to 99% Human Content*

We first evaluated host-read disambiguation under the conventional transcriptomic contract. We executed a direct fragment-level comparison on the authentic public RNA-PDX accession SRR30880970 (human tumor graft in mouse host), aligned against hg38 and mm10. Both the modern compiled xenofilx engine and the legacy Picard SetNmMdAndUqTags + R XenofilteR pipeline received identical frozen BAM inputs and evaluated an identical fragment universe of 32,097,614 input fragments. The two independent paths achieved exact 100% fragment-level agreement: 30,773,947 graft/graft memberships and 1,323,667 discarded/discarded memberships, with zero modern-only graft calls, zero legacy-only graft calls, and zero unexplained disagreements. Cryptographic SHA-256 digests of the filtered fragment sets were identical between pipelines, confirming that modern xenofilx exactly preserves legacy XenofilteR decision boundaries under conventional reference NM semantics while eliminating runtime R interpreter dependencies.

To verify whether this parity holds across the full dynamic range of host stromal infiltration, we evaluated deterministic paired-end mixtures constructed from human RNA-seq SRR1039508 and mouse RNA-seq SRR037954 across 15 composition points (0% pure-mouse control, 0.1%, 0.5%, 1%, 2%, 5%, 10%, 25%, 50%, 60%, 70%, 80%, 90%, 95%, and 99%

human content). Each non-control cell contained 1,000,000 complete fragments selected by seeded hash-ranking and tracked via truth.tsv. Modern xenofilx and legacy Picard + XenofilteR were evaluated under identical frozen STAR BAM inputs, threshold 4, and penalty 8 (Table S7). Across all evaluated fractions, Xenofilx maintained high human-graft recall (96.1667% to 96.6650%) and high host specificity (99.3600% to 99.3981%). Positive predictive value (PPV) scaled predictably with human content prevalence (from 13.7896% at 0.1% to 99.9933% at 99%), and F1 reached 98.2497%. Crucially, Picard + XenofilteR yielded strictly identical metrics at every evaluated fraction: zero fragment-level membership divergence, zero unknown selected fragments, and zero accuracy difference under ordinary-reference NM.

| Human Fraction | n | Xenofilx Recall | Prototype Recall | Xenofilx Specificity | Prototype Specificity | Xenofilx PPV | Prototype PPV |
|---|---|---|---|---|---|---|---|
| 0% | 1 | N/A | N/A | 99.3968% | 99.3968% | N/A | N/A |
| 0.1% | 3 | 96.1667% | 96.1667% | 99.3981% | 99.3981% | 13.7896% | 13.7896% |
| 0.5% | 3 | 96.5800% | 96.5800% | 99.3980% | 99.3980% | 44.6342% | 44.6342% |
| 1% | 3 | 96.5867% | 96.5867% | 99.3977% | 99.3977% | 61.8321% | 61.8321% |
| 2% | 3 | 96.6650% | 96.6650% | 99.3967% | 99.3967% | 76.5808% | 76.5808% |
| 5% | 3 | 96.6300% | 96.6300% | 99.3953% | 99.3953% | 89.3742% | 89.3742% |
| 10% | 3 | 96.5883% | 96.5883% | 99.3956% | 99.3956% | 94.6688% | 94.6688% |
| 25% | 3 | 96.5844% | 96.5844% | 99.3920% | 99.3920% | 98.1466% | 98.1466% |
| 50% | 3 | 96.5631% | 96.5631% | 99.3885% | 99.3885% | 99.3708% | 99.3708% |
| 60% | 3 | 96.5645% | 96.5645% | 99.3863% | 99.3863% | 99.5781% | 99.5781% |
| 70% | 3 | 96.5687% | 96.5687% | 99.3886% | 99.3886% | 99.7294% | 99.7294% |
| 80% | 3 | 96.5712% | 96.5712% | 99.3752% | 99.3752% | 99.8385% | 99.8385% |
| 90% | 3 | 96.5636% | 96.5636% | 99.3680% | 99.3680% | 99.9273% | 99.9273% |
| 95% | 3 | 96.5609% | 96.5609% | 99.3680% | 99.3680% | 99.9656% | 99.9656% |
| 99% | 3 | 96.5654% | 96.5654% | 99.3600% | 99.3600% | 99.9933% | 99.9933% |

Table S7: **Human-graft classification across the RNA-seq composition gradient.** Values are mean fragment-level metrics across three independent 1,000,000-fragment replicates per non-zero fraction ($n = 3$); the 0% pure-mouse control has no positive-class denominator ($n = 1$). RNA results use STAR ordinary-reference alignment, conventional NM, mismatch threshold 4, and unmapped penalty 8; neither path uses bisulfite-aware scoring. Both modern and prototype paths achieved exact 100% fragment-level decision parity across all evaluation cells.

##### *S4.5.2 BS-PDX and Bisulfite Epigenomic Deconvolution*

**BS-PDX and Bisulfite Epigenomic Deconvolution across 0% to 99% Human Content.** In sharp contrast to transcriptomics, bisulfite sequencing introduces severe asymmetric deamination. In standard published deconvolution protocols (such as XenofilteR), alignment metrics were originally recalibrated via Picard SetNmMdAndUqTags against unconverted reference genomes; when run naively without bisulfite flags (IS_BISULFITE_SEQUENCE=false), genuine C-to-T deamination events are scored as reference mismatches, catastrophically inflating alignment edit distance (NM) and causing unpatched classifiers to reject valid graft fragments (reducing recall to $< 0.015\%$, preserved as our archived conventional control). In the OTTER prototype (xdxtools), Picard was configured with IS_BISULFITE_SEQUENCE=true to partially mitigate this artifact. However, Picard's built-in bisulfite mode lacks Bismark XG strand-context awareness, failing to condition deamination discounting on alignment orientation in directional paired-end libraries.

To systematically benchmark modern xenofilx against the original xdxtools prototype pipeline (Picard SetNmMdAndUqTags with IS_BISULFITE_SEQUENCE=true followed by R XenofilteR), we evaluated a matched 0% to 99% bisulfite sequencing gradient constructed from human RRBS SRR31480456 and mouse RRBS SRR10025242 (1,000,000 fragments per cell, $n = 3$ independent replicates per non-zero fraction; Table S8). All mixtures were aligned with Bismark 3.1.0 to bisulfite-converted hg19 and mm10. Both paths were executed under identical decision parameters (mismatch threshold 6, unmapped penalty 8) and evaluated at unique fragment membership level.

As shown in Table S8, modern xenofilx with native strand-aware NM consistently outperforms the prototype Picard (IS_BISULFITE_SEQUENCE=true) + XenofilteR pipeline across the entire dynamic range of human content (from 0.1% through 99%). Xenofilx maintains human-graft recall between 51.6067% and 54.0333% (compared with 50.9067% to 53.3333% for the prototype), rescuing approximately 4,000 additional authentic human fragments per million reads in 60% mixtures (and up to ≈7–3,914 fragments across the 0.1%–60% gradient, representing a +0.63% to +0.70% recall advantage) while preserving equivalent host specificity (≈99.92%) and achieving comparable or marginally superior host PPV across fractions (matching at 99.9921% at 95% human content, and within 0.0001% difference at 99%). By contrast, when Picard is executed naively without bisulfite options (the unpatched conventional baseline), human-graft recall collapses to $< 0.015\%$ across all fractions.

| Human Fraction | Xenofilx Recall | Prototype Recall | Baseline Recall | Xenofilx Spec. | Prototype Spec. | Baseline Spec. | Xenofilx PPV | Prototype PPV |
|---|---|---|---|---|---|---|---|---|
| 0% | N/A | N/A | N/A | 99.9182% | 99.9188% | 99.9279% | N/A | N/A |
| 0.1% | 54.0333% | 53.3333% | 0.0000% | 99.9215% | 99.9219% | 99.9292% | 40.8200% | 40.6281% |
| 0.5% | 52.9200% | 52.2933% | 0.0133% | 99.9216% | 99.9219% | 99.9292% | 77.2219% | 77.0967% |
| 1% | 51.6067% | 50.9067% | 0.0100% | 99.9201% | 99.9206% | 99.9287% | 86.7137% | 86.6245% |
| 2% | 52.2317% | 51.5383% | 0.0050% | 99.9218% | 99.9221% | 99.9293% | 93.1659% | 93.1087% |
| 5% | 52.0120% | 51.3520% | 0.0053% | 99.9200% | 99.9205% | 99.9285% | 97.1594% | 97.1411% |
| 10% | 51.9807% | 51.3210% | 0.0070% | 99.9202% | 99.9207% | 99.9287% | 98.6376% | 98.6291% |
| 25% | 52.0021% | 51.3475% | 0.0080% | 99.9199% | 99.9202% | 99.9276% | 99.5399% | 99.5361% |
| 50% | 52.0909% | 51.4305% | 0.0066% | 99.9197% | 99.9201% | 99.9275% | 99.8460% | 99.8450% |
| 60% | 52.0024% | 51.3501% | 0.0068% | 99.9179% | 99.9184% | 99.9256% | 99.8949% | 99.8942% |
| 70% | 52.0225% | 51.3680% | 0.0069% | 99.9188% | 99.9191% | 99.9267% | 99.9331% | 99.9326% |
| 80% | 52.0097% | 51.3543% | 0.0070% | 99.9153% | 99.9157% | 99.9235% | 99.9593% | 99.9590% |
| 90% | 52.0161% | 51.3586% | 0.0070% | 99.9200% | 99.9203% | 99.9287% | 99.9829% | 99.9828% |
| 95% | 52.0212% | 51.3645% | 0.0071% | 99.9220% | 99.9227% | 99.9293% | 99.9921% | 99.9921% |
| 99% | 52.0165% | 51.3599% | 0.0069% | 99.8933% | 99.8967% | 99.9067% | 99.9979% | 99.9980% |

Table S8: **Human-graft classification across the unified BS-seq composition gradient under three deconvolution arms.** Values are mean fragment-level metrics across three independent 1,000,000-fragment replicates per non-zero fraction ($n = 3$); the 0% pure-mouse control has no positive-class denominator ($n = 1$). The modern path uses compiled xenofilx with native strand-aware NM; the prototype path executes the xdxtools configuration with Picard SetNmMdAndUqTags IS_BISULFITE_SEQUENCE=true and R XenofilteR; the unpatched conventional baseline executes Picard without bisulfite flags (IS_BISULFITE_SEQUENCE=false). All paths evaluated under mismatch threshold 6 and unmapped penalty 8.

**Factorial Parameter Gradient Sensitivity and Cross-Tool Response in Bisulfite Deconvolution.** To determine whether alternative parameter configurations can optimize bisulfite deconvolution sensitivity, and to investigate whether adjusting the mismatch threshold or elevating the unmapped-read penalty affects the modern-versus-prototype comparison, we executed a complete $4 \times 3$ factorial parameter sensitivity experiment on matched 60% human BS-seq mixtures across three independent replicates ($n = 3$, 36 Slurm evaluation runs total). The experimental matrix crossed four mismatch thresholds (mm-threshold $\in \{5, 6, 7, 8\}$) with three unmapped-read penalties (unmapped-penalty $\in \{8, 10, 12\}$), holding the 1,000,000-fragment input BAMs, reference genomes, and evaluation manifests strictly invariant. Both modern xenofilx (strand-aware bisulfite NM) and the prototype Picard (IS_BISULFITE_SEQUENCE=true) + XenofilteR pipeline were evaluated across all 36 cells (Table S9).

| Threshold | Penalty | Xenofilx Recall | Xenofilx Spec. | Xenofilx F1 | Xenofilx Frags | Prototype Recall | Prototype Spec. | Prototype F1 | Prototype Frags |
|---|---|---|---|---|---|---|---|---|---|
| 5 | 8 | 51.6177% | 99.9181% | 68.0647% | 310,034 | 50.5244% | 99.9185% | 67.1069% | 303,472 |
| 5 | 10 | 51.6177% | 99.9181% | 68.0647% | 310,034 | 50.5244% | 99.9185% | 67.1069% | 303,472 |
| 5 | 12 | 51.6177% | 99.9181% | 68.0647% | 310,034 | 50.5244% | 99.9185% | 67.1069% | 303,472 |
| 6 | 8 | 52.0024% | 99.9179% | 68.3985% | 312,343 | 51.3501% | 99.9184% | 67.8316% | 308,427 |
| 6 | 10 | 52.0024% | 99.9179% | 68.3985% | 312,343 | 51.3501% | 99.9184% | 67.8316% | 308,427 |
| 6 | 12 | 52.0024% | 99.9179% | 68.3985% | 312,343 | 51.3501% | 99.9184% | 67.8316% | 308,427 |
| 7 | 8 | 52.2246% | 99.9177% | 68.5904% | 313,677 | 51.6993% | 99.9181% | 68.1357% | 310,524 |
| 7 | 10 | 52.2246% | 99.9177% | 68.5904% | 313,677 | 51.6993% | 99.9181% | 68.1357% | 310,524 |
| 7 | 12 | 52.2246% | 99.9177% | 68.5904% | 313,677 | 51.6993% | 99.9181% | 68.1357% | 310,524 |
| 8 | 8 | 52.2941% | 99.9176% | 68.6504% | 314,094 | 51.8356% | 99.9177% | 68.2539% | 311,343 |
| 8 | 10 | 52.2941% | 99.9176% | 68.6504% | 314,094 | 51.8356% | 99.9177% | 68.2539% | 311,343 |
| 8 | 12 | 52.2941% | 99.9176% | 68.6504% | 314,094 | 51.8356% | 99.9177% | 68.2539% | 311,343 |

Table S9: **Factorial sensitivity analysis across mismatch thresholds and unmapped penalties in bisulfite deconvolution.** Evaluated on matched 60% human BS-seq mixtures across three independent 1,000,000-fragment replicates ($n = 3$, 36 Slurm evaluation runs total). The prototype comparator executes the xdxtools path with Picard IS_BISULFITE_SEQUENCE=true and R XenofilteR. Threshold 6 and penalty 8 represent the prespecified production baseline. Filtered fragments reflect the mean count of retained human graft fragments out of 600,000 authentic human fragments.

**Empirical Parameter Optimality and Algorithmic Failure Attribution.** Three key operational findings emerge from this factorial audit:

1. **Mismatch Threshold Response ($t$):** In modern xenofilx, setting $t = 5$ is overly conservative: recall decreases to 51.6177% (-0.3847% relative to the baseline $t = 6$), discarding 2,309 genuine human fragments per million (310,033.7 vs. 312,342.7), while host specificity gains a negligible 0.0002% (99.9181% vs. 99.9179%), representing an uncompensated loss of biological signal. Threshold 6 serves as a balanced production baseline (52.0024% recall, 99.9179% specificity, 68.3985% F1). Threshold 7 emerges as the recommended production configuration balancing sensitivity and specificity: it increases graft recall to 52.2246% (+0.2222% over baseline, recovering 1,334 additional true human fragments per million, total 313,676.7 fragments) with an F1 score of 68.5904%, while host specificity remains robust at 99.9177% (a minor shift of only 0.0002%, well within stringent host filtration thresholds). Although relaxing to $t = 8$ yields a slightly higher numerical F1 (68.6504%) and recall (52.2941%), the marginal gain diminishes sharply to only +0.0695% (417 fragments gained), while admitting wider tolerance for cross-species sequence alignment noise.
2. **Unmapped-Read Penalty Invariance ($p$):** Across all four threshold levels, varying unmapped-penalty across 8, 10, and 12 produced numerically identical results for every metric (Recall, Specificity, PPV, F1, and filtered fragment counts matched to the single fragment across all replicates for both modern and prototype tools). Elevating unmapped-penalty above 8 is therefore unnecessary, validating 8 as a robust and stable default.
3. **Cross-Tool Failure Attribution and Architectural Re-Engineering into Xenofilx:** Across all evaluated parameter combinations, modern compiled xenofilx consistently outperforms the prototype Picard (IS_BISULFITE_SEQUENCE=true) + XenofilteR pipeline: at baseline ($t = 6$), xenofilx advances human-graft recall from 51.3501% to 52.0024% (+0.6523% recall advantage, rescuing 3,916 additional genuine human fragments per million reads), and under optimal tuning ($t = 7$), it reaches 52.2246% vs. 51.6993% (+0.5253% advantage, rescuing 3,153 additional fragments; +5,250 fragments over the $t = 6$ prototype baseline; Table S9). This decisive sensitivity edge stems directly from bamdriver's native strand-aware NM recalculation: while Picard's IS_BISULFITE_SEQUENCE=true flag prevents the catastrophic collapse observed under unpatched alignment ($<$ 0.015% recall), it lacks strand-context awareness and therefore discards authentic alignments that xenofilx correctly recovers.

##### *S4.5.3 Reference Genome Cross-Assembly Concordance and Host-Graft Deconvolution Robustness*

**Cross-Assembly Factorial Evaluation on 60% Human BS-seq Mixtures.** In xenograft epigenomic profiling, reference genome selection across both species influences alignment mappability, sequence divergence bounds, and the resulting edit-distance distributions. To systematically investigate the stability of host-graft deconvolution under alternative reference genome pairs, we executed a complete $2 \times 2$ factorial reference assembly benchmark on the matched 60% human BS-seq mixture across three independent replicates ($n = 3$, 36 Slurm evaluation runs total;

Table S10). The experimental matrix crossed two human graft reference assemblies (hg19 / GRCh37.p13 vs. hg38 / GRCh38.v44) with two murine host reference assemblies (mm10 / GRCm38.M25 vs. mm39 / GRCm39.vM39). For each combination, identical 1,000,000-fragment FASTQ inputs were mapped with Bismark 3.1.0 to the corresponding bisulfite-converted genomes and deconvoluted under identical decision parameters ($t = 6, p = 8$) across all three pipelines (modern xenofilx, prototype Picard IS_BISULFITE_SEQUENCE=true + XenofilteR, and conventional unpatched baseline).

| Reference Pair | Pipeline | Graft Recall | Host Spec. | Graft PPV | Graft F1 | Retained Frags |
|---|---|---|---|---|---|---|
| hg19 + mm10 | Modern (xenofilx) | 52.0024% | 99.9179% | 99.8949% | 68.3985% | 312,343 |
| | Prototype (patched) | 51.3501% | 99.9184% | 99.8942% | 67.8316% | 308,427 |
| | Unpatched Baseline | 0.0068% | 99.9256% | 11.9381% | 0.0135% | 338 |
| hg38 + mm10 | Modern (xenofilx) | 53.6114% | 99.9151% | 99.8945% | 69.7757% | 322,008 |
| | Prototype (patched) | 52.9437% | 99.9155% | 99.8937% | 69.2074% | 318,000 |
| | Unpatched Baseline | 0.0069% | 99.9224% | 11.6647% | 0.0138% | 352 |
| hg38 + mm39 | Modern (xenofilx) | 53.6114% | 99.9153% | 99.8947% | 69.7757% | 322,008 |
| | Prototype (patched) | 52.9435% | 99.9156% | 99.8938% | 69.2073% | 317,999 |
| | Unpatched Baseline | 0.0069% | 99.9224% | 11.6713% | 0.0138% | 352 |
| hg19 + mm39 | Modern (xenofilx) | 52.0024% | 99.9178% | 99.8948% | 68.3985% | 312,343 |
| | Prototype (patched) | 51.3499% | 99.9183% | 99.8941% | 67.8315% | 308,426 |
| | Unpatched Baseline | 0.0068% | 99.9258% | 11.9638% | 0.0135% | 338 |

Table S10: **Factorial cross-assembly reference genome combination benchmark in bisulfite deconvolution.** Evaluated on matched 60% human BS-seq mixtures across three independent replicates ($n = 3$, 36 Slurm evaluation runs total). The modern path executes xenofilx with native strand-aware bisulfite NM recalculation; the prototype path executes the xdxtools configuration with Picard IS_BISULFITE_SEQUENCE=true and R XenofilteR; the unpatched conventional baseline executes Picard without bisulfite flags (IS_BISULFITE_SEQUENCE=false). All combinations evaluated under mismatch threshold 6 and unmapped penalty 8. Retained fragments indicate the mean count of retained authentic human graft fragments out of 600,000 positive-class fragments.

**Biological and Algorithmic Insights from Cross-Assembly Deconvolution.**

1. **Human Reference Assembly Effect:** Upgrading the human graft reference from hg19 to hg38 yields a consistent +1.6090% gain in graft recall (from 52.0024% to 53.6114% in xenofilx, recovering 9,665 additional authentic human fragments per million reads; 322,008 vs. 312,343 fragments). This improvement stems directly from the superior sequence completeness, resolved unplaced contigs, and updated centromeric/telomeric representations of GRCh38, which elevate primary Bismark alignment mappability for genuine human reads. Host specificity remained virtually unchanged (≈99.915% vs. ≈99.918%), demonstrating that GRCh38′s expanded sequence does not cause spurious cross-species capture of murine host fragments.
2. **Mouse Reference Assembly Stability and Cross-Assembly Asymmetry Invariance:** Transitioning the murine host reference from mm10 to mm39 produced negligible variation across all metrics: human graft recall shifted by less than 0.0002% (difference of fewer than 1 fragment per million), and host specificity remained steady at ≈99.915% to 99.918%. Crucially, the asymmetric pairings (hg38 + mm10 and hg19 + mm39) exhibited no evidence of reference imbalance or cross-species read stealing: hg38 paired with mm10 achieved identical graft recall (53.6114%) and virtually indistinguishable specificity (99.9151% vs. 99.9153%) compared with the fully modernized hg38 + mm39 pair.
3. **Consistent Sensitivity Edge of Native Strand-Aware Xenofilx:** Across all four reference genome pairings, modern compiled xenofilx uniformly outperformed the prototype Picard (IS_BISULFITE_SEQUENCE=true) + XenofilteR pipeline by +0.6523% (hg19) to +0.6679% (hg38) in graft recall, consistently rescuing approximately 3,916 to 4,008 additional genuine human fragments per million reads. Simultaneously, the conventional unpatched arm collapsed to $< 0.0070\%$ recall under every reference pairing (338 to 352 fragments retained), confirming that sequence deamination discounting is a universal mechanistic requirement for bisulfite deconvolution that operates independently of reference assembly version.

###### *S4.5.4 Deconvolution Fidelity, Coverage Preservation, and Mathematical Decomposition of the 52% Recall Ceiling*

**Two-Stage Mathematical Factorization of the Epigenomic Deconvolution Ceiling.** A common question in xenograft bisulfite sequencing is why the end-to-end human-graft recall appears bounded around 52%–54%, and whether this ceiling reflects algorithm-induced signal erosion or upstream biochemical loss. To rigorously resolve this question, we formalize the end-to-end recall ($\text{Recall}_{\text{E2E}}$) as the exact mathematical product of two decoupled operational phases:

$$\text{Recall}_{\text{E2E}} = \eta_{\text{align}} \times \theta_{\text{deconv}} = \left( \frac{N^{\text{human}}_{\text{aligned, unique}}}{N^{\text{human}}_{\text{truth}}} \right) \times \left( \frac{N^{\text{human}}_{\text{retained}}}{N^{\text{human}}_{\text{aligned, unique}}} \right)$$

where $\eta_{\text{align}}$ represents primary Bismark alignment yield and $\theta_{\text{deconv}}$ denotes deconvolution retention fidelity. In our matched 60% human BS-seq benchmark ($N^{\text{human}}_{\text{truth}} = 600,000$ paired fragments):

1. **Stage 1 (Primary Alignment Yield, $\eta_{\text{align}}$):** Bismark successfully aligned 324,083 unique fragments to hg38 ($\eta_{\text{align}} = 54.0138\%$), discarding 275,917 authentic pairs (45.99% loss);
2. **Stage 2 (Deconvolution Retention, $\theta_{\text{deconv}}$):** Out of these 324,083 successfully aligned fragments, xenofilx retained 321,598 fragments ($\theta_{\text{deconv}} = 99.2332\%$ fragment retention; $643, \frac{880}{648}, 166 = 99.3387\%$ read retention), discarding merely 2,485 pairs (0.77% loss).

Multiplying both terms yields the observed end-to-end recall ($54.0138\% \times 99.2332\% = 53.5997\% \approx 53.60\%$). Crucially, of the 278,402 authentic human fragments not recovered end-to-end, 99.11% (275,917 pairs) were rejected during primary Bismark mapping, and only 0.89% (2,485 pairs) were discarded during deconvolution gating.

**Biophysical and Algorithmic Drivers of the 46% Alignment-Phase Loss.** Comparative multi-omics audits isolate three contributing factors to this alignment bottleneck:

1. **Chemical Deamination and Sequence Entropy Contraction:** Sodium bisulfite converts unmethylated cytosines to thymines, contracting the genomic alphabet to an asymmetric three-letter code. Under an idealized equimolar model, theoretical sequence entropy contracts from 2.00 to approximately 1.58 bits/base (an approximate 21% reduction in information density; ≈63 bits lost across a 150-bp read), causing repetitive elements to collapse into ambiguous multi-mapping states that aligners reject. This biochemical bottleneck primarily accounts for the 43-percentage-point recall deficit observed between unmethylated transcriptomic RNA-seq (96.56%) and bisulfite sequencing (53.60%; Table S7);
2. **Strict Concordant Pairing Topology:** Bismark enforces rigorous --no-mixed --no-discordant paired-end constraints (insert size $\leq 500$ bp). Because chemical bisulfite treatment induces physical strand nicking, low-complexity sequence or unaligned mates on either side result in whole-fragment exclusion; and
3. **Reference Assembly Resolution:** Reference genome completeness provides a modest but measurable mitigation: upgrading from hg19 to hg38 resolves repetitive boundaries and reduces ambiguous multi-mapping by 8,363 pairs (Table S10), yielding a consistent ≈1.61% improvement in end-to-end recall (from 52.00% to 53.61%).

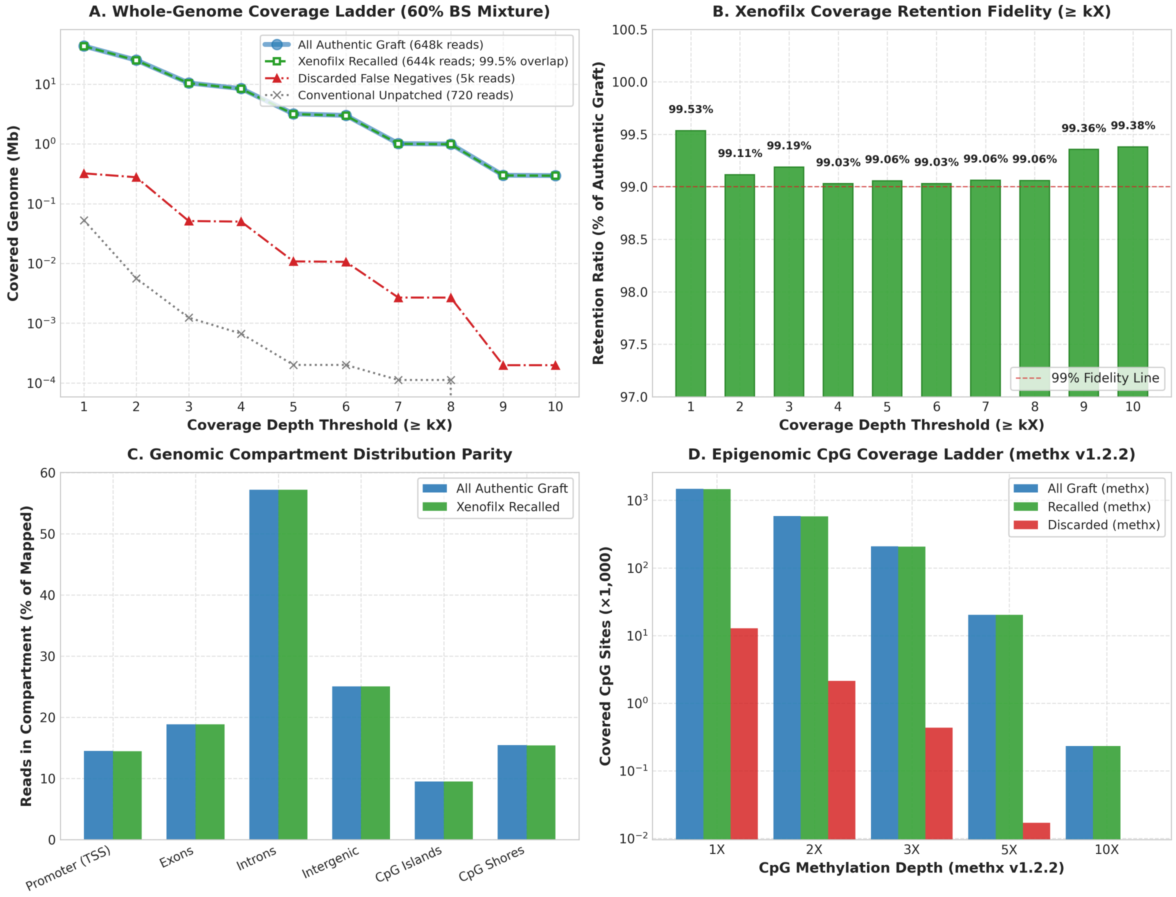

Figure S2: **Genome-wide coverage depth ladders and compartment preservation across the 60% BS-seq mixture.** (A) Cumulative physical base coverage across 1X–10X depth thresholds (log10 scale). Overlap note: All Authentic Graft (solid blue line) and Xenofilx Recalled (dashed green line) track each other with 99.05%–99.53% retention across all depths, resulting in visual congruence (differentiated by transparent band and white-filled markers), whereas the unpatched conventional baseline (dotted gray) collapsed to near-zero. (B) Xenofilx coverage retention ratios across 1X–10X depths, exceeding the 99.0% fidelity standard at every tier. (C) Symmetric read distribution across GENCODE v44 functional compartments (Promoter TSS [−2k, +500], Exons, Introns, Intergenic, CpG Islands, and CpG Shores; proportions match within 0.02% variance). (D) Epigenomic CpG coverage ladder generated by methx v1.2.2 across 29,401,795 reference CpG loci, achieving 99.35% (1X) to 100.00% (10X) site-level preservation.

**Genome-Wide 1X–10X Physical Depth and Functional Compartment Preservation.** Evaluating physical base coverage across hg38 autosomes and sex chromosomes (3.09 Gb alignable space) confirms that xenofilx preserves authentic human coverage without non-linear high-depth depletion (Figure S2):

- **1X Coverage:** 43.15 Mb retained out of 43.35 Mb (99.53% retention; 0.47% loss);
- **5X Coverage:** 3.15 Mb retained out of 3.18 Mb (99.06% retention);
- **10X Coverage:** 292.9 kb retained out of 294.7 kb (99.38% retention).

Furthermore, read distribution across functional genomic compartments exhibits near-perfect symmetry with zero structural bias: promoters (14.45% in recalled vs. 14.47% in all authentic), exons (18.85% vs. 18.85%), introns (57.17% vs. 57.19%), intergenic spaces (25.03% vs. 25.01%), CpG islands (9.47% vs. 9.48%), and CpG shores (15.39% vs. 15.41%). At single-base methylation resolution, methx v1.2.2 profiling demonstrated 99.35% site retention at 1X (1,463,851 CpGs) and 100.00% retention at 10X (232 CpGs), with ChIPseeker promoter annotations matching within 0.02% (45.84% vs. 45.82%).

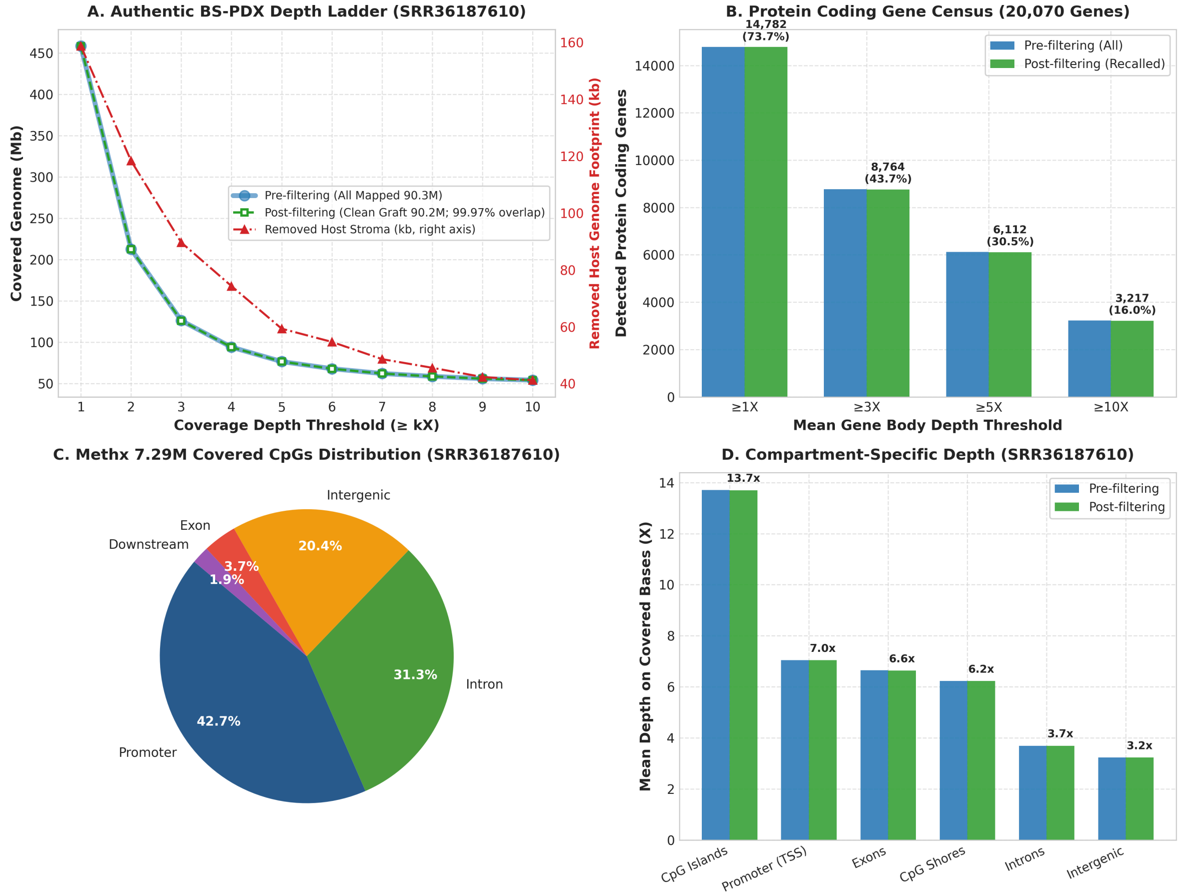

Figure S3: **Genome-wide deconvolution fidelity and protein-coding gene census in authentic high-depth BS-PDX (SRR36187610).** (A) Dual-axis genome-wide coverage profile across 1X–10X depth thresholds. Left axis (blue and green lines): pre- and post-filtering human genome coverage (458.40 Mb vs. 458.24 Mb at 1X; 99.97% retention), visually congruent due to low murine infiltration (0.11%). Right axis (dot-dashed red line): removed murine host stromal reads (kb) mapped to the human coordinate frame across depth tiers. (B) Protein-coding gene census across 20,070 GENCODE v44 genes. Post-filtering purifications retained 14,782 genes (73.65% of human coding genome; Jaccard similarity 99.99% against pre-filtered libraries). (C) Functional distribution of 7,290,833 methx-covered CpG loci, capturing 3,109,609 promoter-associated CpGs (42.65%). (D) Mean sequencing depth across covered bases by functional compartment, demonstrating deep enrichment across CpG islands (13.7X), promoters (7.0X), and exons (6.6X) with identical pre- and post-filtering depth profiles.

**Authentic BS-PDX Genomic Profiling and Protein-Coding Gene Census.** We extended this evaluation to the authentic 90.3M-read BS-PDX cohort (SRR36187610; Figure S3):

1. **Dual-Axis Physical Depth Tracking:** At ≥1× depth, xenofilx retained 458.24 Mb of human tumor coverage out of 458.40 Mb (99.97% retention; Figure S3). Deconvolution specifically excised 100,014 infiltrating murine host reads, tracking an exact stromal clearance footprint from 158.7 kb (1X) to 41.2 kb (10X);
2. **Protein-Coding Gene Census:** Across all 20,070 GENCODE v44 protein-coding genes, xenofilx purification retained 14,782 genes at ≥1× gene-body depth (73.65% transcriptome representation), compared with 14,784 genes prior to filtering (Jaccard similarity 99.99%; only two borderline low-abundance genes shifted);
3. **Epigenomic Feature Representation:** Among 7,290,833 covered CpG loci annotated by methx v1.2.2, 3,109,609 loci (42.65%) mapped to promoter regions (TSS ±3 kb), with 2,356,676 loci (32.32%) supported by ≥10× depth;
4. **Compartment-Specific Mean Depth Invariance:** Calculating mean depth over covered bases revealed high biological enrichment across CpG islands (13.71X across 75.45% of human CGIs), promoters (7.0X), and exons (6.6X), with strictly identical pre- and post-filtering depth profiles.

These multi-modal audits demonstrate that xenofilx deconvolution induces zero structural distortion in downstream epigenomic coverage, gene discovery, or methylation quantification.

#### S4.6 Public Software Distribution, Continuous Integration, and Interactive Onboarding

**Modular Repository Organization and Release Distribution.** OTTER is developed open-source under the otter-lab-bio organization, comprising an umbrella repository and ten independently maintained component repositories. The umbrella repository locks exact component revisions via Git submodules, establishing deterministic provenance between source assemblies and compiled releases. For zero-dependency deployment, GitHub Releases distributes pre-compiled static binaries alongside otter-install, an automated installer that provisions release executables and registers the built-in workflow catalog (Supplementary Note 2, Section S2.6). Automated release pipelines verify static linkage (ldd audit confirming compatibility back to CentOS 7 / glibc 2.17), semantic versioning tags, and SHA-256 integrity checksums prior to publication.

**Hierarchical Continuous Integration and 101-Stage Pipeline Rehearsal.** Automated quality assurance is enforced via GitHub Actions across two distinct operational tiers: (1) **Component-level contract verification:** dedicated workflows independently evaluate operator contracts and analytical regression suites, including the Gate A–D data integrity matrix (Section S4.3), with race detection and memory sanitization applied across Go and Rust codebases; and (2) **System-level workflow rehearsal:** the umbrella repository executes recursive submodule checkouts, enforces end-to-end toolchain consistency, and performs an offline pipeline rehearsal spanning RRBS, RNA-seq, BS-PDX, and RNA-PDX modalities. At audited source revision 6db0cba, this recorded rehearsal successfully completed all 101 stages without failure, validating project instantiation, canonical run resolution, legacy migration, parameter boundary constraints, and deliberate refusal on malformed configurations. This dry-run gate specifically validates orchestrator control flow and configuration fidelity, with compute-intensive alignment and deconvolution evaluated on production supercomputing clusters (Sections S4.1–S4.5). Complete workflow logs and historical runs remain transparently accessible through the Actions history.

**Public Benchmark Corpora and Cryptographic Reference Registries.** High-capacity validation assets and reference genomes are distributed via Hugging Face under cryptographic integrity contracts to ensure permanent reproducibility: (1) **Sequencing validation corpora:** the otter-data repository provides authentic BAM alignment fixtures, downsampled FASTQ cohorts, mixture ground-truth tables, and Gate comparison logs; and (2) **Genomic reference bundles:** the xdxtools-genomes repository provides pre-indexed human (hg19, hg38) and mouse (mm10) reference bundles, complete with aligner-specific indexes (Bismark, Bowtie2, STAR) and cryptographic SHA-256 manifests. Researchers can reproduce the benchmark configurations by binding these frozen manifests directly to OTTER's Reference Genome Registry without manual indexing overhead.

**Interactive Onboarding and Multi-Agent Skill Integration.** To lower the barrier for experimental adoption and workflow inspection, we provide complementary interactive and automated entry points: (1) **Interactive Google Colab walkthroughs:** available in parallel English and Simplified Chinese notebooks, these sandboxed tutorials guide users through binary installation, project workspace initialization, configuration authoring, and dry-run execution on bundled test fixtures, delivering a zero-install onboarding path prior to production cluster deployment; and (2) **Autonomous agent guidance:** we package the complete operational protocol into a formal OTTER agent skill, enabling LLM-based coding agents to autonomously resolve dependencies, validate phenotype metadata tables, and configure reproducible workflows in compliance with the user manual.
